# Proprotein convertase activity regulates cumulus-oocyte-complex matrix integrity and cumulus cell migration during ovulation via a GDF9-dependent mechanism

**DOI:** 10.64898/2026.07.22.738304

**Authors:** Caroline E. Kratka, Ruixu Huang, Jeffrey Pea, Robin M. Skory, Christina D. King, Jiyang Zhang, Pawat Pattarawat, Caroline M. Milner, Anthony J. Day, Nicolas Plachta, Shuo Xiao, Birgit Schilling, Darryl L. Russell, Brittany A. Goods, Francesca E. Duncan

## Abstract

Cumulus cells have well-established roles early in ovulation but the key molecules that drive their behavior in later stages, leading to follicle rupture, remain underexplored. Here, we observed that inhibition of proprotein convertases (PCSKs) via a pan-inhibitor (PCI) impaired follicular rupture and disrupted the cumulus matrix integrity within intact follicles. Reduced cumulus cell adherence to the cumulus-oocyte-complex (COC) matrix was also observed in isolated COCs and notably occurred late during the maturation window without affecting oocyte maturation. Visualization of PCSK transcript and protein expression, as well as selective inhibition of specific PCSKs, determined that the observed phenotype in COCs is likely attributed to PCSK5A inhibition. We conducted bulk RNA-sequencing and proteomics of PCI-treated COCs which revealed that PCSK inhibition caused dysregulation of extracellular matrix organization, cell migration/adhesion, and TGF-β signaling pathways. Subsequent validation showed that this inhibition translated to disrupted matrix organization and altered migratory and adhesive behaviors in cumulus cells. The TGF-β ligand GDF9 has a predicted PCSK cleavage site, and supplementation with GDF9 rescued matrix integrity suggesting its role as a downstream substrate of PCSKs to regulate matrix organization. Altogether, this study identified PCSK5A and GDF9 as key regulators of COC matrix integrity and cumulus cell migration during late ovulation. These findings highlight novel factors required for follicle rupture which can be leveraged for the development of fertility therapeutics and contraceptives.

## Introduction

Ovulation, the process by which an egg is released from the ovary for potential fertilization by sperm, occurs hundreds of times throughout a woman’s lifetime and is critical for both ovarian and systemic health^1–10^. Prior to ovulation, the ovary produces large amounts of estrogen, which has extraovarian benefits on the musculoskeletal, cardiovascular, and central nervous systems^11–21^. The ovary also produces progesterone as a result of ovulation and subsequent luteinization, which supports pregnancy in response to successful fertilization or continued cycling in its absence. Disorders impacting ovulation, such as polyendocrine metabolic ovarian syndrome, are leading causes of infertility^22,23^ (Teede et al., 2026). Therefore, an understanding of the factors regulating ovulation has important clinical implications for fertility, contraception, and overall women’s health.

Several key events with distinct regulatory mechanisms must occur during ovulation, including oocyte meiotic maturation, cumulus-oocyte-complex (COC) expansion, follicle wall thinning at the apex, and follicular rupture. Critical to the success of these events is the activity of cumulus cells, which are specialized granulosa cells that surround the oocyte within antral follicles. Cumulus cells provide the oocyte with metabolic support, regulate meiotic maturation, and improve oocyte quality^24–30^. Following the initiation of ovulation via the luteinizing hormone (LH) surge, LH stimulates the downstream effectors EGFR, ERK1/2, C/EBPα/β, and PGR in granulosa cells^31,32^. These regulators drive production of factors important for expansion of the extracellular matrix (ECM) around the COC, such as TSG-6, PTX3, and PTGS2^32–35^. COC expansion is thought to contribute to ovulation and fertilization, as loss of matrix regulators and subsequent disruptions in matrix formation can severely impair fertility^36–40^.

Recent ‘omics studies of the ovary during ovulation have expanded the catalogue of molecular players^41,42^. In addition, *ex vivo* models that recapitulate the morphology, endocrinology, and physical dynamics of *in vivo* ovulation have enabled identification of novel molecular signatures of mammalian ovulation^43–48^. Analysis of a time course of *ex vivo* ovulation using single follicle transcriptomics revealed a group of genes whose expression is continuously upregulated during ovulation, suggesting an active role in this process^47^. Of note, several members of the proprotein convertase family, including *Pcsk3*, *Pcsk5*, and *Pcsk6*, exhibit this specific pattern of expression. The proprotein convertases (PCSKs) are serine endoproteases which are widely expressed and have different substrates^49^. In the ovary, conditional knockout of *Pcsk3* in the oocytes of primordial (*Gdf9*-Cre) and primary (*Zp3*-Cre) follicles leads to early secondary follicle arrest and thereby failure of spontaneous ovulation^50^. Treatment of antral follicles *ex vivo* with a pan-PCSK inhibitor (proprotein convertase inhibitor (PCI)) blocks follicle rupture^47^. However, the mechanism of PCSK action in the follicle during ovulation has not been defined.

In this study, PCSKs were further investigated to determine their role in cumulus cell function during ovulation. Beyond blocked follicle rupture, PCSK inhibition via PCI led to complete disruption of cumulus cell matrix integrity in intact follicles. This phenotype was also observed when COCs were treated directly with PCI during *in vitro* maturation (IVM). PCSK*3* and *5a,* relative to PCSK*5b* and *6,* exhibited enriched transcript and protein expression in the COC that increased during *in vivo* ovulation. However, treatment of COCs with a PCSK3-specific inhibitor did not impact the cumulus matrix, demonstrating that the observed phenotype was likely solely due to PCSK5A inhibition. To determine the mechanism by which PCI treatment causes dissolution of the cumulus matrix, we performed transcriptomic and proteomic analyses. There was striking concordance between the COC transcriptome and proteome, which both revealed dysregulation of processes related to ECM organization, cell adhesion, migration, and metabolism. These molecular signatures translated into changes in cumulus cell function, including matrix disorganization, altered migratory and adhesion behavior, and increased cell blebbing. While PCSKs have multiple substrates, the oocyte secreted factor and member of the TGF-β superfamily, GDF9, contains a potential cleavage site, and PCI treatment of the COC caused dysregulation of the TGF-β pathway. In fact, supplementation with recombinant GDF9 rescued the disruption of COC matrix integrity induced by PCI. These findings provide evidence of a novel ovulatory mechanism by which cumulus cell-derived PCSK activity regulates GDF9 to coordinate the precise timing of matrix integrity and dissolution combined with cumulus cell migratory behavior required for the highly orchestrated process of ovulation.

## Results

### PCSK inhibition blocks follicular rupture in an ex vivo model of ovulation via effects on the COC

We previously reported that the expression of several *Pcsk* family members increases throughout the time course of ovulation *in vitro* and *in vivo*, and that blocking the activity of these proteins via pharmacologic inhibition prevents follicular rupture in an *ex vivo* model of ovulation^47^. However, it is unknown whether this rupture failure is due solely to effects on the follicular wall or whether PCI also impacts the COC. Therefore, we performed *ex vivo* ovulation in the presence or absence of 10 μM PCI in intact follicles as previously described^47^ and examined the morphology of the COC (**Fig 1A**). PCI is a reversible competitive substrate analog that is a pan-inhibitor of the PCSKs (Ki = 16 pM)^51^. As observed previously, follicles cultured under control conditions ruptured, whereas PCI-treated follicles did not (**Fig 1B**)^47^. In contrast to fully expanded COCs released from control follicles, eggs were nearly completely denuded of matrix and cumulus cells when micro-dissected from unruptured PCI-treated follicles (**Fig 1B**). Consistent with this, PCI treatment was associated with a decrease in *Has2* expression in follicles at 4 h post-hCG relative to controls (32 ± 2 a.u. vs. 14 ± 3 a.u. in CTL vs. PCI-treated follicles, respectively; p<0.0001) (**Fig 1C)**. *Has2* is the primary hyaluronan synthase expressed in cumulus cells that mediates cumulus expansion during ovulation^38,52^. Thus, these results indicate that PCSK inhibition disrupts the COC matrix, which may influence follicular rupture given that the perturbation of cumulus cells is linked to failed ovulation^36–40^.

**Figure 1:**
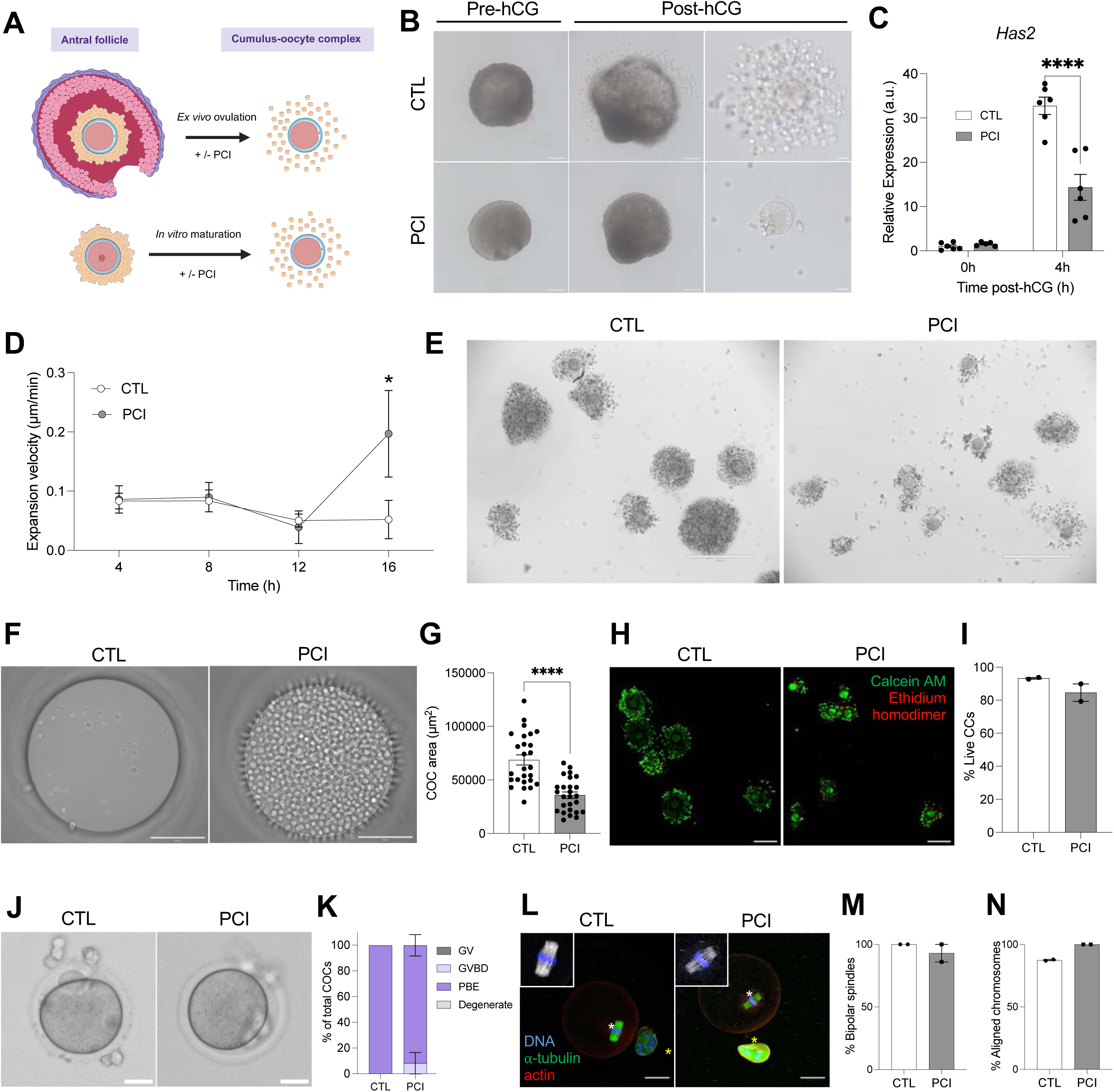
PCSK inhibition blocks follicular rupture in an ex vivo model of ovulation via effects on the COC. A) Schematic showing methodologies for testing the direct effect of PCI on antral follicles (top) and COCs (bottom). B) Left: Time-lapse images from videos of follicles cultured *ex vivo* with or without PCI. Scale bar = 100 µm. Right: Brightfield images of COCs removed from control or PCI-treated follicles. Scale bar = 25 µm. C) *Has2* expression in follicles treated with PCI for 4 h. Data are represented as mean ± SEM. **** p < 0.0001 by two-way ANOVA. n=5-6 follicles per treatment group. D) Expansion velocity of CTL and PCI-treated COCs across in vitro maturation in the EmbryoScope+^TM^. Data are represented as mean ± SEM. * p < 0.05 by mixed-effects analysis. n=48 biological replicates per treatment group. E) Representative brightfield images of COCs treated with PCI or DMSO control (CTL) following culture and transfer out of the EmbryoScope+^TM^ dish. F) Representative brightfield images of residual cumulus cells in EmbryoScope+^TM^ wells after removal of CTL or PCI-treated COCs. G) Graph showing COC area of CTL or PCI-treated COCs following transfer out of the EmbryoScope+^TM^ dish. Data are represented as mean ± SEM. **** p < 0.0001 by Welch’s t-test. n=26 biological replicates per treatment group. H) Live/dead assay of COCs cultured in CTL or PCI-treated media for 16 h in the standard incubator. Live cells are shown in green (calcein AM) and dead cells are shown in red (ethidium homodimer). Scale bar = 200 µm. I) Quantification of live cumulus cells relative to total cumulus cells in CTL and PCI-treated COCs. Data are represented as mean ± SEM. p > 0.05 by Fisher’s exact test. n=2 biological replicates (5-14 pooled COCs per treatment group). J) Representative brightfield images of CTL vs. PCI treated COCs after 16 h culture and denuding. K) Graph showing maturation status of CTL or PCI-treated COCs after 16 h of culture. Data are represented as mean ± SEM. p>0.05 by Fisher’s exact test. n=15-16 biological replicates per treatment group. L) Immunofluorescent images depicting chromosomes (DAPI; blue), spindles (alpha-tubulin; green), and actin (rhodamine phalloidin; red) in CTL or PCI-treated COCs after denuding of cumulus cells. White asterisks denote spindles and yellow asterisks denote the polar body. Scale bar is 25 µm. Graphs showing percent of M) COCs with bipolar spindles and N) aligned chromosomes after culture with or without PCI. For M) and N), Data are represented as mean ± SEM, p>0.05 by Fisher’s exact test, and n=3 biological replicates (15-16 pooled COCs per treatment group).

To confirm that PCSK inhibition was specific to cumulus cells, we determined the impact of PCI on isolated COCs via IVM using time-lapse microscopy (**Fig 1A, Sup Video 1**). PCI-treated COCs exhibited similar expansion dynamics early in the culture period relative to controls. However, beginning at 12 h of culture, cumulus cells from PCI-treated COCs appeared to detach from the matrix and rapidly migrated to the edges of the culture well (**Sup Video 1**). This phenotype translated into increased expansion velocity after 12 h (0.05 µm/min ± 0.03 vs. 0.2 ± 0.07 µm/min in CTL vs. 10 µM PCI-treated COCs at 16 h, respectively; p=0.04), further confirming the effect of PCSK inhibition late in the maturation process (**Fig 1D**)^53^. This phenotype occurred in a dose-dependent manner (**Fig 1A**, **Sup Fig 1, Sup Video 1**), and 10 μM PCI was used thereafter to remain consistent with the paradigm used in the intact follicle^47^. When COCs were transferred out of the microwells following IVM, PCI-treated COCs were almost fully denuded, compared to control COCs which had intact expanded matrices. The loss of an intact matrix with PCSK inhibition suggests complete dissolution of the matrix and detachment of cumulus cells from the complex (**Fig 1E**). This was further demonstrated in wells in which control COCs had been cultured which retained few residual cumulus cells after transfer, whereas those from PCI-treated COCs contained many detached cumulus cells, consistent with loss of cumulus cell adherence to the matrix (**Fig 1F**). As a result, COCs treated with PCI exhibited a smaller final COC area relative to controls (p<0.0001) (**Fig 1G**). This phenotype was not due to cell toxicity as there was no difference in the percentage of live cumulus cells between control (93 ± 0.0055 %) and PCI-treated COCs (85 ± 0.053 %) (p = 0.60) and all oocytes were viable (**Fig 1H-I**).

While PCSK inhibition affected cumulus cells, this phenotype was independent of effects on the oocyte. When oocytes were isolated from *in vitro* matured COCs, similar percentages of cells reached the mature metaphase II (MII)-arrested state irrespective of PCI treatment (100 ± 0% vs. 92 ± 8.3% for control and PCI-treated COCs respectively, p=0.29) (**Fig 1J-K**). The MII eggs derived from PCI treatment exhibited normal bipolar spindles (93 ± 7.0% vs. 100 ± 0.0% COCs with bipolar spindles in CTL vs. PCI-treated COCs, respectively; p=0.48) with aligned chromosomes on the metaphase plate (94 ± 6.0% vs. 94 ± 6.5% COCs with chromosomes aligned at the metaphase plate in CTL vs. PCI-treated COCs, respectively; p>0.99) (**Fig 1L-N**). Moreover, when we performed *in vitro* fertilization (IVF) on COCs exposed to PCI during IVM, fertilization and embryonic development rates were similar to *in vitro* matured controls, supporting that PCSK inhibition does not affect the developmental competence of the egg (**Sup Fig 2, Sup Video 2**).

We further validated that PCSK inhibition did not impact the oocyte directly by performing spontaneous IVM on denuded oocytes in the presence or absence of PCI (**Sup Fig 3**, **Sup Video 3**). Timelapse microscopy revealed similar morphokinetic parameters of meiotic progression, including time to germinal vesicle breakdown (0.95 ± 0.029 h vs. 1.0 ± 0.042 h, p=0.28), time to polar body extrusion (8.1 ± 0.096 h vs. 8.4 ± 0.073 h in control vs. PCI respectively, p=0.051), and duration of meiosis I (7.2 h ± 0.098 vs. 7.4 h ± 0.059 in control vs. PCI respectively, p=0.11) (**Sup Fig 3A-B**). Similar percentages of cells in the PCI-treated and control groups reached the MII stage (97 ± 2.1% vs. 100 ± 0% for control and PCI-treated COCs respectively, p=0.4226) (**Sup Fig 3C**), and PCI treatment did not affect either bipolar spindle formation in the resulting eggs (100 ± 0.0% vs. 100 ± 0.0% COCs with bipolar spindles in CTL vs. PCI-treated COCs, respectively; p>0.99) or chromosome alignment (88 ± 3.2% vs. 83 ± 8.3% COCs with aligned chromosomes at the metaphase plate in CTL vs. PCI-treated COCs, respectively; p=0.21) (**Sup Fig D-F**). Taken together, these findings provide strong evidence that the observed effect of PCSK inhibition is specific to cumulus cells.

### Pcsk5a is the primary Pcsk regulating cumulus cell function during ovulation

Given that PCI is a pan-PCSK inhibitor and that three members of the PCSK family (*Pcsk3*, *5*, and *6*) were previously shown to be actively upregulated in follicles during ovulation^47^, we wanted to better understand which PCSK was likely responsible for the observed phenotype. First, we mined the expression profiles of *Pcsk3*, *5*, and *6* across ovarian cell types in a single-cell transcriptomic dataset of the mouse ovary throughout ovulation (**Fig 2A**)^41^. This dataset identified two distinct cumulus cell clusters: Cumulus 1 and Cumulus 2 enriched at 4 h post-hCG (early) and at 12 h post-hCG (late), respectively. *Pcsk5* was highly expressed in both early and late cumulus cell clusters, whereas *Pcsk3* and *6* had lower expression. We further validated these expression patterns via RNA *in situ* hybridization using histological sections of mouse ovaries across a time course of ovulation (**Fig 2B, Sup Fig 4**). This assay also allowed us to distinguish between the isoforms: *Pcsk5a* and *Pcsk5b*. *Pcsk5a* is widely expressed (including in the ovary) whereas *Pcsk5b* is primarily expressed in the small intestine, kidney, and liver^47,54,55^. Previously, we observed that *Pcsk5a* relative to *Psck5b* was enriched in the whole follicle and induced by hCG^47^. Ovaries were analyzed at 0 h, 4 h, 8 h, and 11 h post-hCG, corresponding to early, mid, and late stages of the ovulatory process, respectively^41,42,47,56^. Consistent with previous findings, we observed little expression of *Pcsk5b* in the COC and whole ovary across ovulation (**Fig 2B-C, Sup Fig 4**). We found that hCG induced minimal expression of *Pcsk6* in the COC and mural granulosa cells, but *Pcsk6* was primarily expressed in preantral follicles (**Fig 2B-C, Sup Fig 4**). On the contrary, *Pcsk3 and 5a* were both highly expressed in the COC (**Fig 2B-C, Sup Fig 4B-C**). The normalized cumulus cell expression of both *Pcsk3* and *Pcsk5a* peaked at 11 h post-hCG (0.84 ± 0.025 a.u. and 0.81 ± 0.014 a.u., respectively) out of all timepoints tested and was significantly greater than *Pcsk5b* or *Pcsk6* at this timepoint (0.053 ± 0.0093 a.u. and 0.89 ± 0.022 a.u., respectively; p<0.0001) (**Fig 2B-C, Sup Fig 4B-C)**. Although the expression patterns of *Pcsk3* and *5a* were similar throughout ovulation (p<0.05 at all timepoints), *Pcsk3* was also detected in the oocyte, whereas *Pcsk5a* expression was limited to cumulus cells (**Fig 2B, Sup Fig 4B**). Based on these results, we concluded that *Pcsk3* and *5a* are present in the COC and thus more likely to regulate cumulus cell function, as opposed to *Pcsk5b* or *6*.

**Figure 2:**
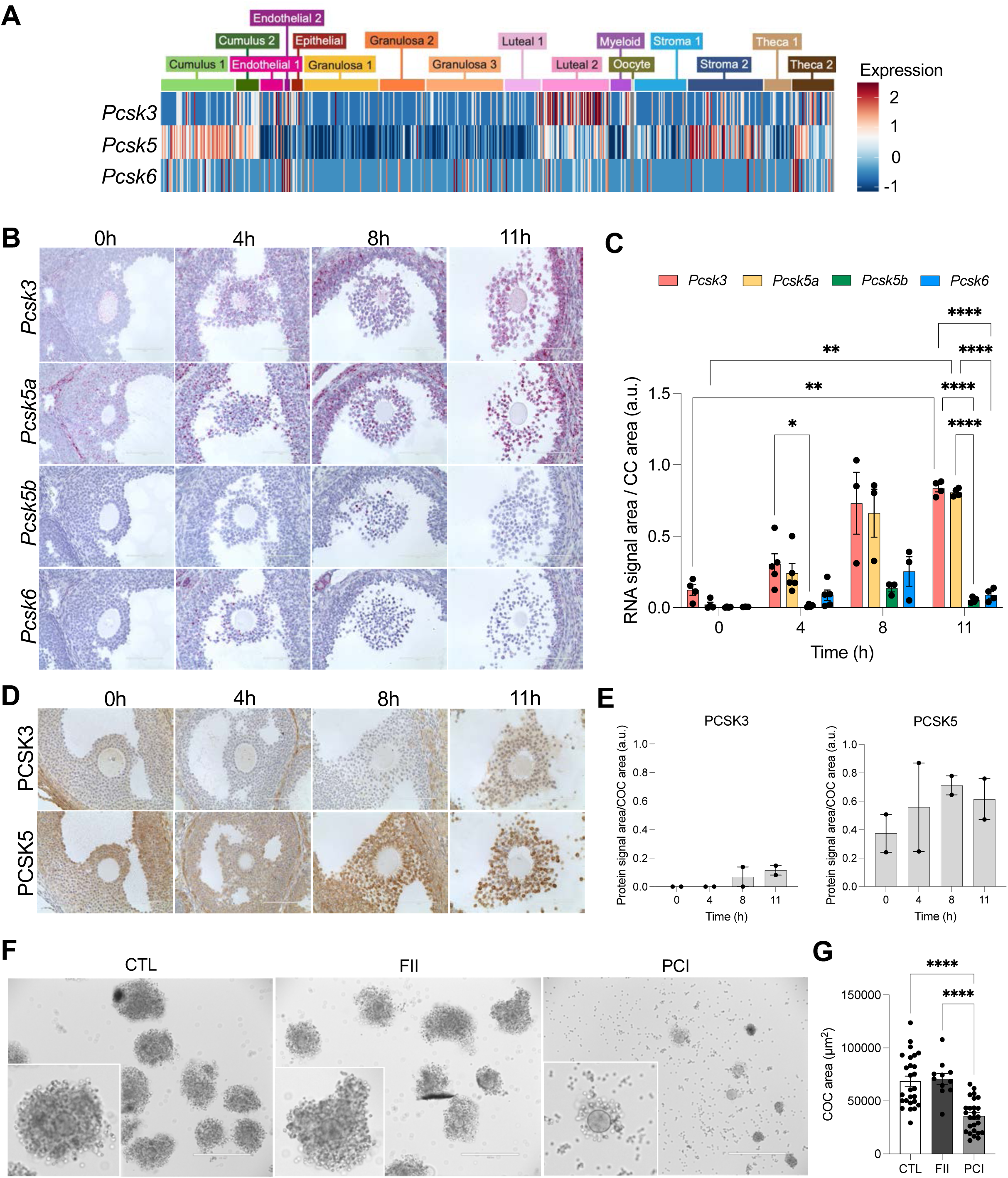
PCSK5A is the primary PCSK regulating cumulus cell function during ovulation. A) Heatmap depicting expression of *Pcsk3*, *5*, and *6* in early (4 h post-hCG) and late (12 h post-hCG) cumulus cells from a published single cell dataset of the mouse ovary throughout an *in vivo* time course of ovulation^41^. B) Representative brightfield images from RNA *in situ* hybridization depicting expression of *Pcsk3* and *Pcsk5* in COCs at 0 h, 4 h, 8 h, or 11 h post-hCG injection. C) Quantification of *Pcsk3*, *5*, and *6* expressions throughout ovulation. Data are represented as mean ± SEM. p-values as follows: * < 0.05, ** < 0.01, *** < 0.001, **** < 0.0001 by mixed-effects analysis. n=3-5 biological replicates per *Pcsk*. D) Representative brightfield images from immunohistochemistry depicting abundance of *Pcsk3* and *Pcsk5* in COCs at 0 h, 4 h, 8 h, or 11 h post-hCG injection. E) Quantification of PCSK3 and PCSK5 protein throughout ovulation. Data are represented as mean ± SEM. p>0.05 for all timepoint comparisons by one-way ANOVA. n=2 biological replicates per PCSK. F) Representative brightfield images of COCs treated with proprotein convertase inhibitor (PCI), Furin inhibitor (FII), or DMSO control (CTL) following culture and transfer out of the EmbryoScope+^TM^ dish. G) Graph showing COC area of CTL, FII-, or PCI-treated COCs following transfer out of the EmbryoScope+^TM^ dish. Data are represented as mean ± SEM. CTL and PCI data replotted from Fig 1E (experiments conducted in parallel). **** p < 0.0001 by Welch’s t-test. n=11-26 biological replicates.

Because protein expression is a more direct reflection of function, we next performed immunohistochemistry on sequential ovarian sections to those used for RNA *in situ* hybridization to better understand PCSK3 and PCSK5 expression dynamics in COCs across the time course of ovulation. PCSK3 expression was low in the COC until 8 h post-hCG, whereas PCSK5 was expressed continuously in the COC throughout ovulation (**Fig 2D-E, Sup Fig 5**). These findings support that PCSK3 and PCSK5A are both expressed in the COC and thus are potential candidates driving PCSK activity. Therefore, to distinguish between PCSK3 and PCSK5 activity, we determined whether a specific inhibitor for PCSK3 (Furin inhibitor II; FII; Hexa-D-Arginine) had a similar effect on the COC as PCI. Unlike PCI treatment which resulted in near complete dissolution of the COC, FII-treated COCs remained fully intact similar to controls (**Fig 2F-G, Sup Fig 6**). These results suggest that PCSK5A, rather than PSCK3, is the primary PSCK regulating the COC during ovulation. Consistent with this, FII was previously tested in parallel to PCI in *ex vivo* ovulation assays and was less effective in inhibiting follicular rupture than PCI^47^.

### PCSK inhibition accelerates the normal temporal pattern of COC gene expression and results in dysregulation of pathways related to the ECM, cell adhesion, cell migration, and metabolism

To elucidate the molecular mechanism by which PCSK inhibition affects matrix integrity, we performed bulk RNA-sequencing of PCI-treated COCs relative to controls at 0 h, 4 h, 8 h, or 12 h of culture during IVM (**Fig 3A**). PCI-treated and control COCs exhibited similar gene expression profiles at 4 h (7 upregulated and 49 downregulated genes in PCI-treated COCs relative to control), consistent with their comparable expansion behavior and morphology at this time point (**Fig 3B-C**, **Fig 1D; Sup Fig 1, Sup Video 1, Sup Files 1 and 2**). By 8 h and 12 h of culture, the transcriptomic differences between PCI-treated COCs and controls were more prominent with 1,613 genes (880 upregulated and 733 downregulated) and 1,327 genes (501 upregulated and 826 downregulated) exhibiting differential expression, respectively (**Fig 3C, Sup Files 1 and 2**). These results indicate that changes in the transcriptome precede the phenotypic effects of PCSK inhibition on the COC matrix which were observed around 12 h (**Fig 3B-C, Sup Fig 1 and 7A, Sup Video 1**). Notably, unbiased clustering revealed that 8 h PCI-treated COCs clustered more closely with 12 h control COCs than 8 h control COCs, suggesting that PCSK inhibition accelerates aspects of the normal temporal expression profile of COCs, rather than resulting in a completely dysregulated expression pattern (**Fig 3B, Sup Fig 7A**).

**Figure 3:**
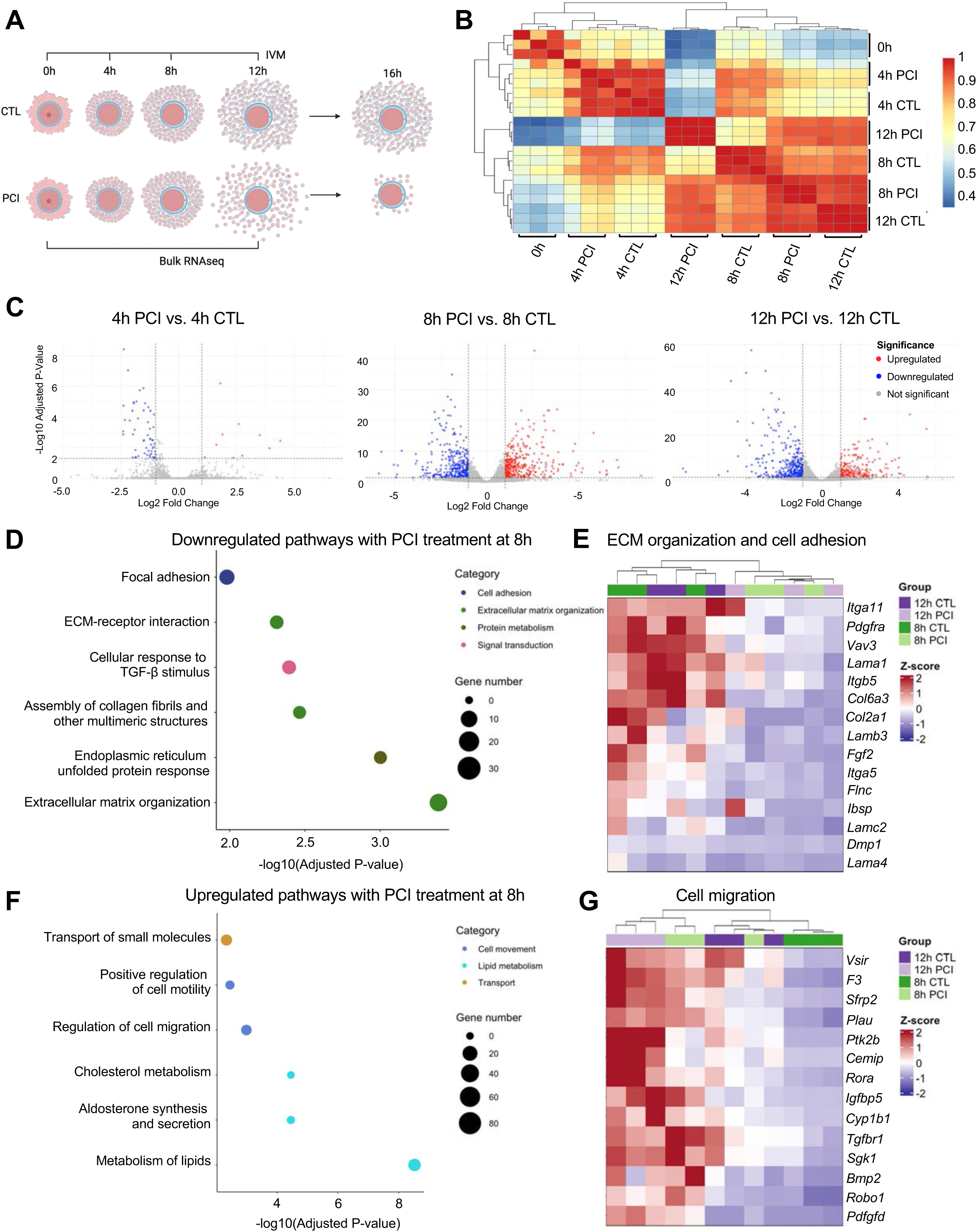
PCSK inhibition accelerates the normal temporal pattern of COC gene expression and results in dysregulation of pathways related to the ECM, cell adhesion, cell migration, and metabolism. A) Schematic showing methodology for collecting COCs for bulk sequencing. IVM = *in vitro* maturation. B) Heatmap showing relationships between bulk sequenced samples with unbiased clustering. Red and blue indicate high and low correlations between samples, respectively. C) Volcano plots showing number of up- and down-regulated genes in PCI-treated COCs relative to control COCs at each timepoint. D) Dot plot showing pathways downregulated with PCI treatment at 8 h. E) Heatmap of top genes associated with ECM organization and cell adhesion that are downregulated by PCI treatment. F) Dot plot showing pathways upregulated with PCI treatment in 8 h COCs. G) Heatmap of top genes associated with cell migration that are upregulated by PCI treatment.

To better understand the potential functions of differentially expressed genes between PCI-treated COCs and controls, we performed gene ontology analysis (**Fig 3D-G, Sup File 3**). We then categorized the pathways by their broad function and extracted the genes driving the pathways in categories of interest (**Sup Table 1**). Pathways predicted to be downregulated with PCI treatment at both 8 h and 12 h of culture were primarily related to the ECM and focal adhesions which is consistent with the associated loss of matrix integrity (**Fig 3D-E**, **Fig 1B and 1E-F; Sup Fig 1, Sup Video 1, Sup File 3, Sup Table 1**). Indeed, among the genes most affected by PCI treatment were ECM components including collagens (*Col6a3*, *Col2a1*), integrins (*Itga11*, *Itgb5*, *Itga5*), and laminins (*Lama1*, *Lamb3*, *Lamc2*, *Lama4*) (**Fig 3E, Sup File 3, Sup Table 1**). Other downregulated pathways included TGF-β signaling (*Cited1*, *Smad7*, *Smad9*, *Dab2*, *Bambi*, *Id1*, *Sox9*, and *Sox5*) and endoplasmic reticulum (ER) stress responses (**Fig 3D, Sup Fig 8A, Sup File 3, Sup Table 1**). Impaired TGF-β signaling is consistent with PCSK inhibition, as TGF-β ligands are established PCSK targets^57–60^.

Pathways related to lipid metabolism, cell migration, and small molecule transport were enriched in PCI-treated COCs at 8 h and 12 h relative to controls (**Fig 3F-G and Sup Fig 8B, Sup File 3, Sup Table 1**). The most differentially expressed metabolic genes had functions in steroidogenesis (*Cyp11a1*), steroid hormone catabolism (*Cyp1b1*), inositol phosphate metabolism (*Itpka*, *Pik3c2g*), oxidative phosphorylation (*Coq8a*), glycolipid transport (*Gm2a*), and hyaluronan turnover (*Cemip*) (**Sup Fig 8C, Sup File 3, Sup Table 1**). Upregulation of genes associated with lipid metabolism is consistent with data demonstrating that cumulus cells at 12 h post-hCG exhibit enrichment of lipid metabolic pathways compared to those at 4 h post-hCG^41^. Cumulus cells are also known to exhibit migratory, invasive, and adhesive behaviors which peak just before ovulation and have been likened to cancer cells^61^. PCI-treated COCs exhibited upregulation of genes associated with motility, such as *Cemip*, *Igfbp5*, and *Ptk2b* at 8h post-hCG (**Fig 3F-G, Sup File 3, Sup Table 1**). Thus, instability of the COC matrix as a result of PCSK inhibition likely creates a permissive environment for cumulus cells to more freely migrate away from the ECM and other cumulus cells. Notably, multiple of the top migration-related genes were also categorized as metabolism-related by Enrichr, including *Cemip*, *Bmp2*, *Vsir*, *Clec7a*, *F3*, *Rora*, *Cyp1b1*, *Robo1*, and *Tgfbr1* (**Fig 3G and Sup Fig 8C, Sup File 3, Sup Table 1**).

Given the close clustering of 8 h PCI-treated COCs with 12 h control COCs, we compared the similarities in gene expression among these groups. Many of the shared enriched pathways were the same pathways that were upregulated with PCI treatment at 8 h relative to controls at the same time, including those related to cell migration and pathways in cancer (**Sup Fig 8D, Sup File 3**). The expression of these genes at 8 h post-hCG with PCI treatment is consistent with an accelerated onset of typical cumulus cell expression patterns which are likely important for the late stages of ovulation as the COC detaches from the follicle wall in preparation for release from the follicle^61^.

### The COC proteome parallels the transcriptome following PCSK-inhibition

To determine whether transcriptomic changes in the COC following PCSK inhibition were reflected at the protein level, we performed proteomics on COCs that were *in vitro* matured in the presence or absence of PCI for 10 h. This timepoint was selected because it is after initiation of major transcriptomic changes (8 h) but prior to when the COC matrix was observed to fall apart with PCI treatment (∼12 h) (**Fig 4A, Sup Fig 1, Sup Video 1**). A total of 5,631 proteins were detected in untreated COCs at this timepoint, and pathway analysis revealed that major biological processes included mRNA translation, metabolism, immune function, and mRNA surveillance (**Sup Fig 9A, Sup File 5**). There were 79 differentially expressed proteins between PCI-treated COCs and controls that distinguished these cohorts (**Fig 4B-C, Sup Fig 7B and 9A, Sup File 6**). This discrete set of differentially expressed proteins suggests a highly targeted response to PCSK inhibition. Eighteen of these differentially expressed proteins met the classification for core matrisome (ECM glycoproteins, and proteoglycans) or matrisome-associated (ECM regulators, ECM-affiliated proteins, and secreted factors) proteins (**Fig 4D and Sup File 7**)^62^. The majority of proteins with dysregulated expression were involved in pathways related to cell adhesion/migration, ECM remodeling, metabolism, signaling, and stress response (**Fig 4E, Sup Fig 9B-C**). Some of the most differentially expressed proteins with PCI treatment included PLAU and CSPG4 (ECM remodeling), IGFBP5 and TNS4 (adhesion/migration), and TGFBR1 and FST (TGF-β signaling) (**Fig 4F, Sup File 6**). Importantly, the expression of several oocyte-specific or enriched proteins was unaffected by PCSK treatment, providing further evidence that the effect of PCI is specific to cumulus cells. **Sup Table 2** contains a list of oocyte-specific and enriched proteins that were stable with PCI treatment. Interestingly, the pathways underlying the proteomic differences were similar to those observed with the transcriptomics analysis (**Fig 4E-F**, **Fig 3D-F, Sup File 3**). In fact, there was tight correlation between transcript and protein expression at the 8 h and 10 h timepoints, respectively (r=0.898, p=4.99E^-22^) (**Fig 4G**). Such high coordination between the COC transcriptome and proteome likely reflects the rapid nature of ovulation, which only spans a matter of hours and thus requires efficient execution of functions necessary for follicle rupture.

**Figure 4:**
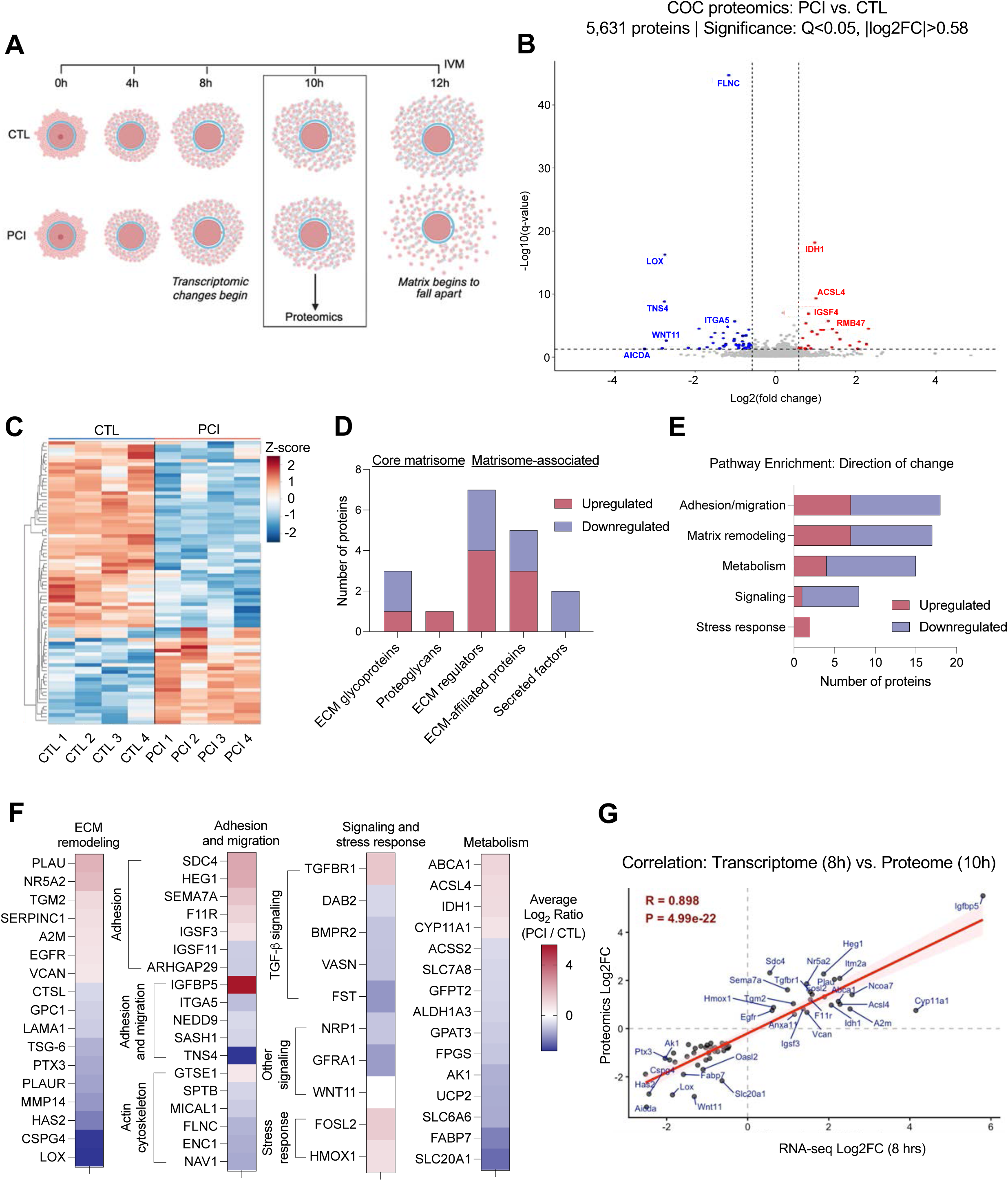
The COC proteome parallels the transcriptome following PCSK-inhibition. A) Schematic showing methodology for collecting COCs for proteomics. IVM = *in vitro* maturation. B) Volcano plot showing proteins up- and down-regulated with PCI treatment of COCs. C) Heatmap showing differential protein expression in CTL vs. PCI-treated COCs. D) Bar graph showing core and matrisome-associated proteins dysregulated with PCI treatment. E) Bar graph showing number of up- and downregulated proteins across functional categories. F) Heatmaps showing specific proteins related to matrix remodeling, adhesion/migration, signaling, or stress responses that are altered with PCI treatment. G) Graph showing correlation between transcriptomic and proteomic data of CTL vs. PCI treated COCs at 8h and 10h of IVM, respectively.

### Molecular changes due to PCSK inhibition translate into phenotypic changes in COC matrix organization

Given that the prominent phenotype of PCSK inhibition was loss of matrix integrity, we further examined the COC ECM. The COC ECM is primarily composed of hyaluronan (HA) and organized into a scaffold containing heavy chains (HC) from the inter-alpha-inhibitor family (covalently attached to HA), PTX3 and VCAN, amongst other proteins (e.g., PLAU/PLAUR) (**Fig 5A**)^36–40,52,63–67^. TSG-6 catalyzes the transfer of HC1, HC2 and HC3 from inter-alpha-inhibitor and pre-alpha-inhibitor onto hyaluronan to form ‘HC•HA’^36,38–40,64,68–72^; PTX3, through its interaction with HCs, plays an essential role to crosslink HC•HA complexes into a matrix^73,74^. HA is also bound directly to the proteoglycan versican^66,67,70^. Each of these components is critical for COC matrix formation^37–40,40,71,72,75^. *Has2* and *Ptx3* transcripts were downregulated in PCI-treated COCs compared to controls beginning at 4 h of culture, and the protein levels showed similar patterns (**Fig 5B-C; Sup Files 1, 2, and 5**). This downregulation of *Has2* expression was similar to what was observed when intact follicles treated with PCI were profiled relative to controls (**Fig 1C**). In contrast, *Vcan* transcripts were upregulated in PCI-treated COCs compared to control at 8 h culture with protein levels also increased (**Fig 5B-C; Sup Files 1, 2, and 5**). PCI treatment was associated with a decrease in TSG-6 protein abundance but not transcript (**Fig 5C, Sup Fig 10A; Sup Files 1, 2, and 5**). In addition to key COC ECM genes, we also observed dysregulation of the protease PLAU and its receptor PLAUR (**Fig 5C and Sup Fig 10A; Sup Files 1, 2, and 5**), which together promote proteolysis and ECM remodeling in the COC^63^. We did not observe changes in the expression or abundance of inter-alpha-inhibitor or pre-alpha-inhibitor components including bikunin or the heavy chains. However, this was expected because inter-alpha-inhibitor and pre-alpha-inhibitor are produced by the liver and present in serum^76^. Thus, exogenous serum in the culture media provides a continuous source of inter- and pre-alpha-inhibitor components.

**Figure 5:**
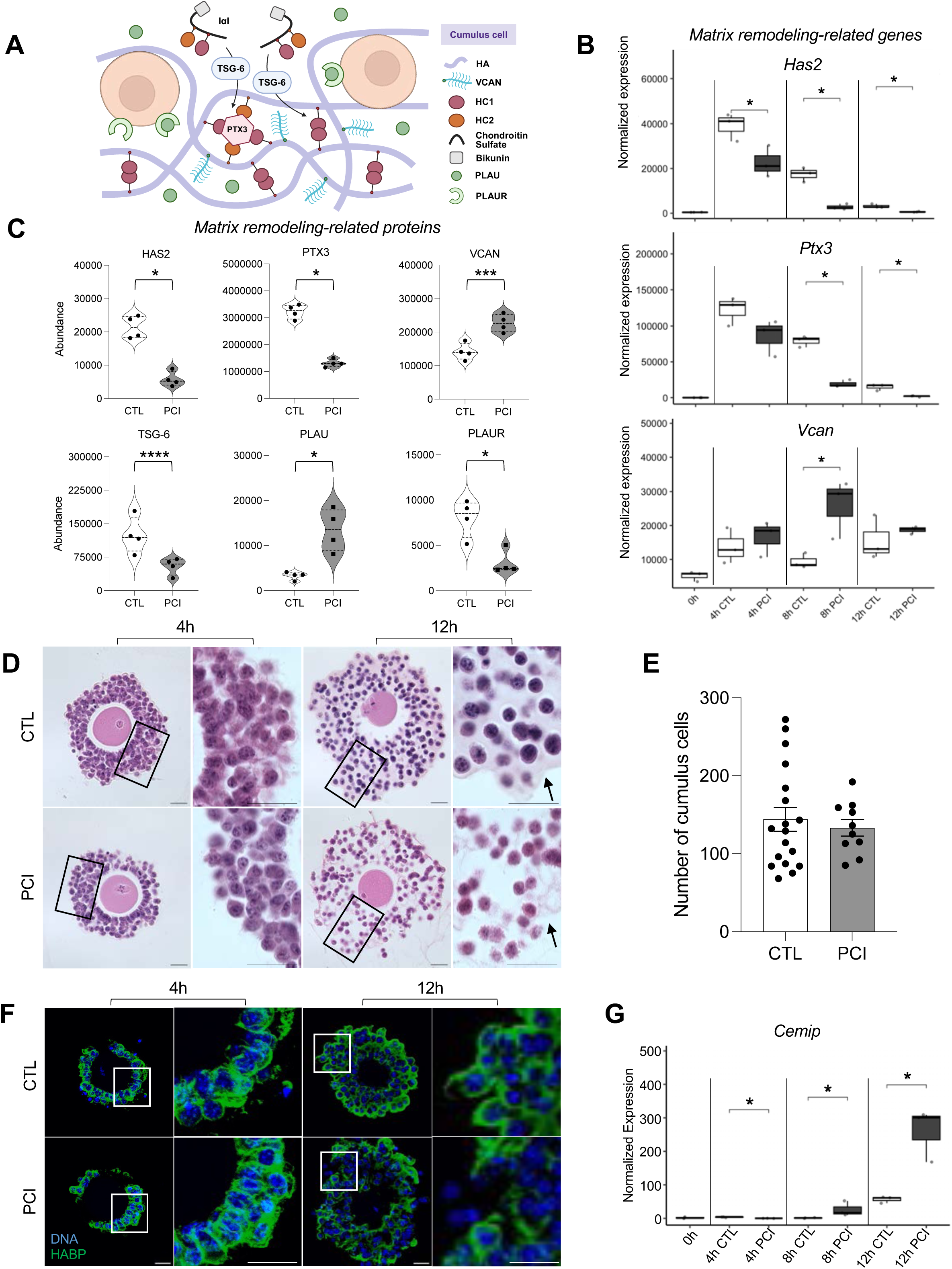
Molecular changes due to PCSK inhibition translate into phenotypic changes in COC matrix organization. A) Schematic showing composition of COC ECM, with components of interest in bold. B) Box and whisker plots showing expression of key COC ECM genes including *Has2*, *Vcan*, and *Ptx3* in CTL vs. PCI treated COCs. Data are represented as mean ± SD. Adjusted p-values for box plots as follows: * < 0.05, ** < 0.01, *** < 0.001, **** < 0.0001 by Wilcoxon signed-rank test. n=3 biological replicates of 10 pooled COCs per treatment and timepoint. C) Violin plots of proteins related to matrix remodeling that are altered with PCI treatment. Data are represented as mean ± SEM. q-values for violin plots as follows: * < 0.05, ** < 0.01, *** < 0.001, **** < 0.0001 by t-test with Storey’s method. n=4 biological replicates of 150 pooled COCs per treatment and timepoint. D) Representative brightfield images of CTL or PCI-treated COCs cultured for 12 in agarose micromolds and stained with hematoxylin and eosin. Boxes highlight locations where high magnification images were captured. Arrows point to edge of ECM in CTL and PCI-treated COCs. E) Graph showing comparison of cumulus cell count between CTL vs. PCI-treated COCs. Data are represented as mean ± SEM. p > 0.05 by Welch’s t-test. n=10-18 COCs across 4 trials. F) Immunofluorescent images of hyaluronan stained with hyaluronic acid binding protein (HABP; green) and chromosomes (DAPI; blue) in control vs. PCI-treated COCs. Scale bar = 25 µm. **** p < 0.0001 by Welch’s t-test. G) Box plot of *Cemip* expression in CTL vs. PCI-treated COCs across *in vitro* maturation. * p < 0.05 by Wilcoxon signed-rank test. n=3 biological replicates of 10 pooled COCs per treatment and timepoint.

The molecular changes associated with PCSK inhibition were reflected in disorganization of the ECM. To probe the architecture of the COC matrix, we performed histology in COCs matured with and without PCI treatment. After 4 h of culture, the organization of the cumulus cells surrounding the oocyte was similar between COCs treated with PCI and controls (**Fig 5D**). However, after 12 h of culture, the PCI-treated COCs exhibited disorganized cumulus cell layers and a loss of the eosin staining in between the cells. Eosin stains proteins non-specifically, so its loss with PCI treatment reflects fewer ECM proteins and/or disrupted matrix organization (**Fig 5D**)^77^. Cumulus cell numbers were similar across cohorts (171 ± 26.7 cells in PCI-treated COCs vs. 169 ± 11.8 cells in control COCs, respectively; p=0.955) demonstrating that cell number differences do not underlie the PCI phenotype (**Fig 5E)**. Given that HA synthesized by HAS2 is a major component of the COC matrix and that HAS2 transcript and protein levels were downregulated with PCI treatment, we visualized HA in COCs matured in the presence or absence of PCI. Although the COC matrix appeared morphologically similar at 4 h, overall HA intensity was lower in PCI-treated COCs relative to controls at this timepoint (0.42 ± 0.024 a.u. vs. 0.31 ± 0.028 a.u. in CTL vs. PCI-treated COCs, respectively; p=0.048), consistent with the reduction in *Has2* transcript and protein observed with PCI treatment of follicles and COCs (**Fig 5F**, **Fig 5B-C, and Fig 1C; Sup Fig 11A**). In contrast, HA intensity was similar between CTL and PCI-treated COCs at 12 h (0.40 ± 0.029 a.u. vs. 0.38 ± 0.051 in CTL vs. PCI-treated COCs, respectively; p=0.97) but the HA matrix exhibited disorganization following PCI treatment relative to controls (**Fig 5F, Sup Fig 11**). In these COCs, multiple cumulus cells lacked surrounding HA, and there were large gaps in the matrix (**Fig 5F**). Interestingly, *Cemip*, a protein involved in HA degradation, was upregulated with PCI treatment consistent with failure to maintain an assembled and organized HA matrix (**Fig 5G**, **Fig 1C and 1E-F; Sup Fig 1, Sup Video 1, Sup Files 1 and 2**). These results together demonstrate that PCSK activity functions late in the ovulatory process to regulate ECM organization and matrix integrity. *Loss of COC matrix integrity due to PCSK inhibition is associated with changes in cumulus cell migration behavior and adhesive properties*

The ECM provides a physical scaffold and mechanical signals that guide cell migration^78–85^. At the transcriptome level, key genes related to cell migration, including *Igfbp5*, and *Ptk2b*, exhibited upregulation with PCI treatment beginning at 8 h (**Fig 6A**). *Igfbp5* is a chemoattractant and is expressed in the bovine and human COC^86–88^. *Ptk2b* is a tyrosine kinase with established roles in actin reorganization and cell migration^89–91^. IGFPB5 is also highly upregulated with PCI treatment at the proteome level, whereas PTK2B protein levels were not affected by PCI (**Sup Fig 12A-B**). To determine whether there were differences in cumulus cell migration with PCSK inhibition, we performed 3D time-lapse imaging of SPY-DNA-labelled COCs and tracked individual cumulus cells over 14 h of IVM (**Fig 6B, Sup Video 4**). Mean square displacement analysis revealed anomalous superdiffusion (α = 1.205 ± 0.255, where α is the anomalous diffusion exponent; **Fig 6C**), consistent with directed, matrix-driven cell displacement rather than passive diffusion. To determine whether proteolytic matrix remodeling underlies this directed motion, we treated COCs with PCI prior to and during expansion. MSD analysis demonstrated a significant reduction in the anomalous diffusion exponent in treated COCs compared to controls (treated α = 1.127 ± 0.295 vs. control α = 1.205 ± 0.255, permutation test p<0.0001) (**Fig 6C**). Despite this reduction in directionality, PCI-treated cumulus cells exhibited greater absolute displacement at equivalent lag times (MSD at t = 55 min: 60.8 μm^2^ vs 39.1 μm^2^; mean displacement 7.8 μm and 6.2 μm; **Fig 6D**), suggesting that the mature cumulus matrix promotes coordinated directional movement while limiting overall cell displacement.

**Figure 6:**
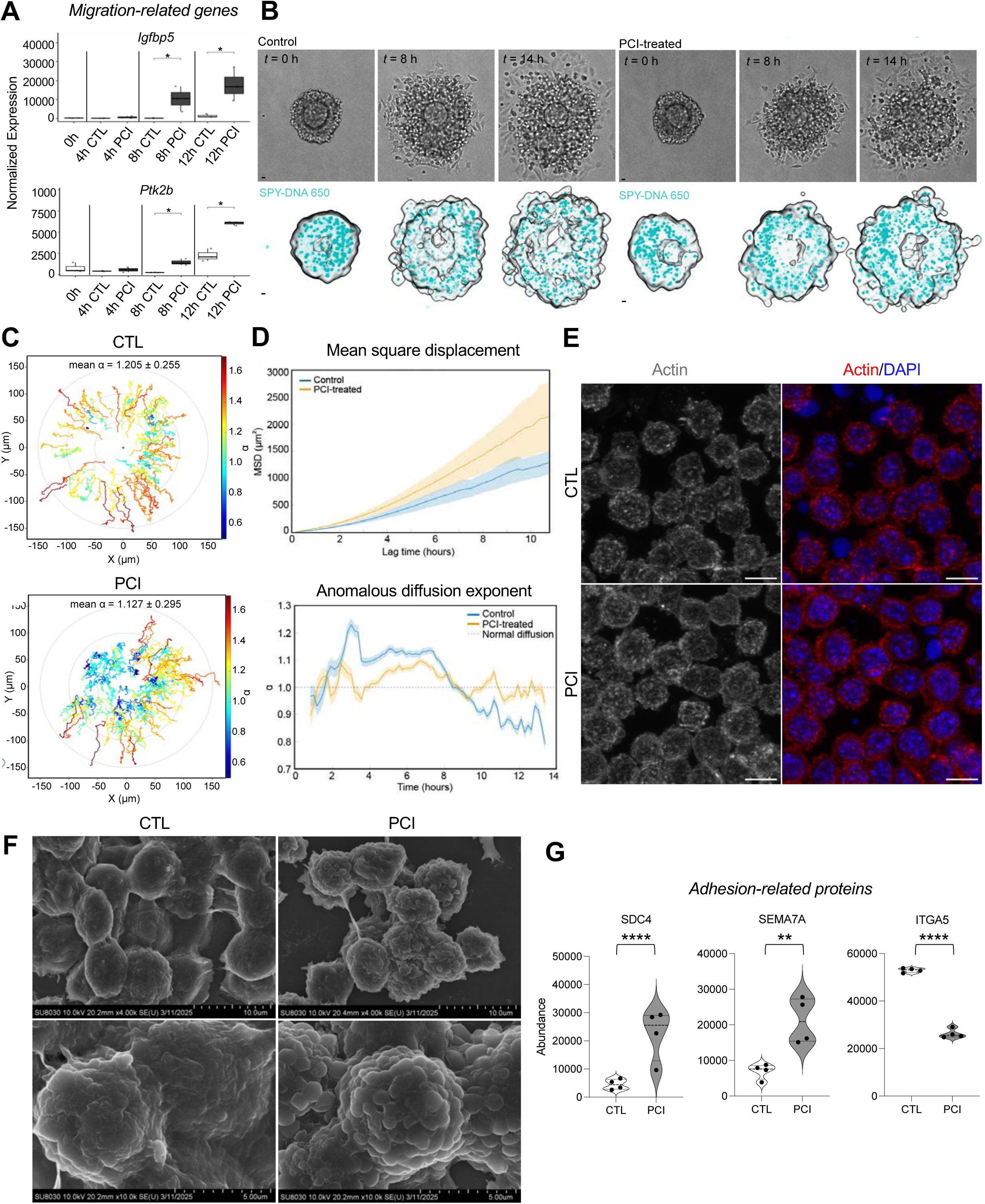
Loss of COC matrix integrity due to PCSK inhibition is associated with changes in cumulus cell migration behavior and adhesive properties. A) Box plots of candidate genes related to migration with are upregulated with PCI treatment. Data are represented as mean ± SD. Adjusted p-values for box plots as follows: * < 0.05, ** < 0.01, *** < 0.001, **** < 0.0001 by Wilcoxon signed-rank test. n=3 biological replicates of 10 pooled COCs per treatment and timepoint. B) Representative brightfield images of CTL COCs stained with SPY-DNA 650 throughout IVM to track cumulus cell migration, with accompanying 3D projections. C) Representative displacement plots of CTL and PCI-treated COCs. Analysis was performed in Imaris (Bitplane) with tracks limited to those with duration greater than 6.5 h. D) Top: Mean square displacement plots of control and PCI-treated COCs, shown with ± SEM. Bottom: Graph showing the anomalous diffusion exponent (L) of CTL vs. PCI-treated COCs relative to normal diffusion (L=1). E) Immunofluorescent images of actin (gray/red) and chromosomes (DAPI; blue) in control vs. PCI-treated COCs. Scale bar = 25 µm. F) Scanning electron microscopy images of CTL and PCI-treated cumulus cells. G) Violin plots of proteins related to cell adhesion that are altered with PCI treatment. Data are represented as mean ± SEM. q-values for violin plots as follows: * < 0.05, ** < 0.01, *** < 0.001, **** < 0.0001 by t-test with Storey’s method. n=4 biological replicates, where each replicate has 150 COCs per treatment group.

To resolve the temporal dynamics of convertase-dependent motility, we performed a sliding window analysis of α using 10-frame windows (∼1.8 h; **Fig 6D**). Both groups exhibited similar behavior during the first ∼2 hours of expansion (p>0.05). Thereafter, control and PCI- treated cumulus cells diverged significantly, remaining divergent through 14 h (p<0.05, per window permutation test; **Fig 6D**). Control cumulus cells exhibited a pronounced peak in α at approximately 2.5 hours (α ≈ 1.23), corresponding to the phase of active matrix remodeling, followed by a sustained superdiffusive plateau (α ≈ 1.12) through ∼8 h. Subsequently, motility transitioned to subdiffusion, consistent with confinement of cells within a mature, crosslinked matrix. In contrast, PCI-treated cumulus cells showed a blunted early peak (α ≈ 1.09), failed to sustain the superdiffusive plateau, and did not undergo the late transition to subdiffusion, indicating impaired matrix maturation and a lack of confinement (**Fig 6D**).

Cumulus cell migration involves remodeling of the actin cytoskeleton into plasma membrane blebs^92^. Actin-enriched blebs were observed in both PCI-treated and control COCs at 12 h of culture (**Fig 6E**). Scanning electron microcopy enabled examination of the topography of the CC surface at high resolution (**Fig 6F**). The blebs were more pronounced in PCI-treated COCs compared to controls which may be related to differential cell motility patterns (**Fig 6B-D, F**). Together, these findings support a model in which PCSK activity drives a temporally regulated program of cumulus cell motility: an initial convertase-independent phase, followed by a convertase-dependent period of directed, matrix-driven expansion, and culminating in a matrix-mediated confinement as the cumulus matures.

Cell migration is intricately linked to cell adhesion, and we noted general gene and protein expression changes in cell adhesion pathways (**Fig 6G**, **Fig 3D-E**, **Fig 4E-F, Sup File 3, Sup Table 1**). Specifically, SDC4 is a heparan sulfate proteoglycan involved in focal adhesion formation^93,94^ which was upregulated in the presence of PCI (**Fig 6G**). However, *Sdc4* transcript was not detected in bulk RNAseq of control and PCI-treated COCs (**Sup Files 1-2**). In addition, SEMA7A, a progesterone receptor-regulated membrane protein expressed in granulosa cells and cumulus cells throughout ovulation is upregulated with PCI treatment (**Fig 6G, Sup Fig 10B, File 6**). SEMA7A is thought to facilitate follicle remodeling via binding to heterodimeric integrins, which can either increase or decrease cell adhesion (referred to as adhesive or repulsive receptors)^95^. One of the adhesive receptors for SEMA7A, integrin α5β1, which is also the primary receptor for fibronectin, is typically upregulated in the COC during ovulation. Here we found that its alpha subunit, ITGA5, was highly downregulated with PCI treatment at the transcriptome and proteome level (**Fig 6G**, **Sup Fig 10B, Sup File 6**)^95^. Loss of ITGA5 (and thus α5β1) is consistent with disruption of focal adhesions in the presence of PCI (**Fig 3D-E**). Given this perturbation of cell adhesion molecules following PCSK inhibition, we examined adhesive properties by culturing mouse COCs at 11 h post-ovulation induction on fibronectin-coated plates for an additional 12 h (**Sup Fig 12C**). The increased adhesion index in COCs after 12 h of PCI treatment (Sup Fig 12D) is consistent with the increased area of cumulus cells in contact with the culture well surface due to loss of matrix integrity, which frees cumulus cells to adhere and migrate over the surface to a greater extent than control cumulus cells.

### PCSK inhibition in the COC is mediated via a GDF9-based mechanism

PCSKs are serine proteases, and each family member has distinct substrate specificities and biological functions. TGF-β ligands were of particular interest because they are known PCSK substrates in other cell types and specifically for PCSK5A^57–60^, which based on our data, is most likely the target of PCI inhibition. Interestingly, treatment of COCs with PCI induced significant alterations in TGF-β related genes and proteins, including TGFBR1, FST, BMPR2, and DAB2 (**Fig 3D, 4E-F, and 7A; Sup Fig 8A and 10C; Sup Files 1, 2, and 5**).

Another critical TGF-β ligand in the context of the COC is GDF9, an oocyte-secreted factor that is required for COC expansion as it promotes expression of COC matrix factors such as *Has2*, *Ptx3*, and *Tsg-6*^96–104^. GDF9 is initially translated as a proprotein and contains a PCSK consensus sequence (**Fig 7B**)^105^. The GDF9 proprotein is known to be cleaved by “furin-like proteases,” but the specific protease has not been identified^106^. Intriguingly, COCs from mice with reduced expression of TGF-β ligands *Bmp15* and *Gdf9* (*Bmp15*^-/-^/*Gdf9*^+/-^) show weak adherence of cumulus cells, similar to the phenotype observed in COCs treated with PCI (**Fig 1E-F; Sup Fig 8A; Sup Fig 1, Sup Video 1**)^96^. Furthermore, genes and/or proteins including *Has2*, *Vcan*, *Cyp11a1*, *Sdc4*, and *Tgfbr1* are dysregulated in COCs when GDF9 signaling is reduced by exogenous heparin or the canonical TGF-β receptor inhibitor SB431542, and these genes are also dysregulated by PCI treatment (**Fig 3D, 4E-F; Sup Fig 8A; Sup Files 1, 2, and 5**)^97^.

**Figure 7:**
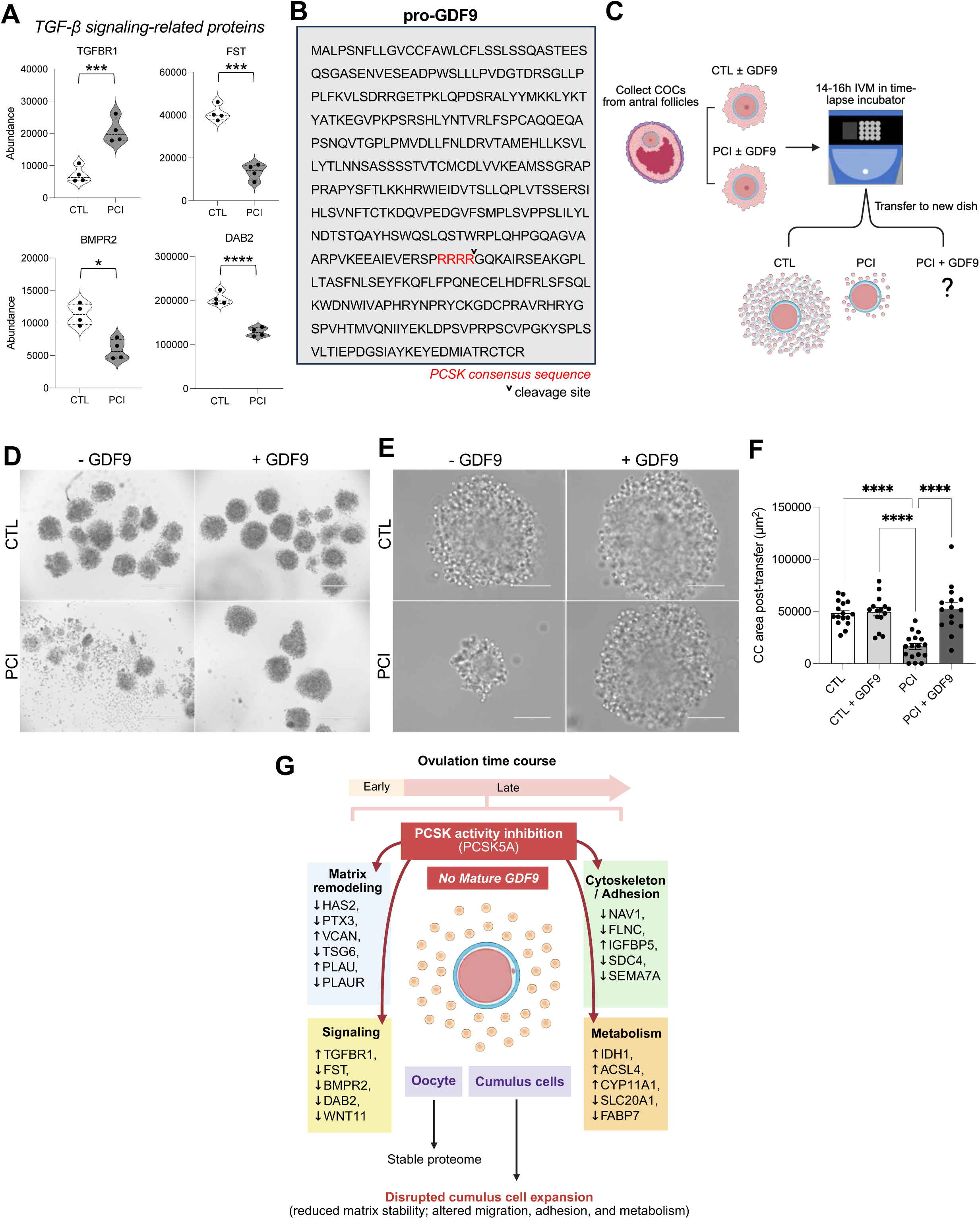
PCSK inhibition in the COC is mediated via a GDF9-based mechanism. A) Violin plots of proteins related to TGF-β signaling that are altered with PCI treatment. Data are represented as mean ± SEM. q-values for violin plots as follows: * < 0.05, ** < 0.01, *** < 0.001, **** < 0.0001 by t-test with Storey’s method. n=4 biological replicates, where each replicate has 150 COCs per treatment group. B) Sequence of pro-GDF9. Red letters highlight the predicted PCSK cleavage site. The down arrowhead indicates where cleavage would occur. C) Schematic showing methodology for GDF9 rescue experiment. D) Brightfield images of COCs cultured for 16h in the EmbryoScope+^TM^ in the presence or absence of PCI and GDF9, followed by transfer to a new culture dish. E) Representative high magnification images of D. F) Graph showing cumulus cell (CC) area of CTL and PCI-treated COCs with or without GDF9 supplementation following culture and transfer. **** p < 0.0001 by one-way ANOVA. G) Schematic summarizing key findings of study.

To determine whether GDF9 may play a role in PCSK-mediated matrix integrity, we matured PCI-treated and control COCs in the presence of recombinant active GDF9 (**Fig 7C**). Remarkably, the addition of GDF9 to PCI-treated COCs completely rescued the PCI phenotype (**Fig 7D-F, Sup Video 5**). When the COCs were transferred at the conclusion of culture, the PCI + GDF9-treated COCs remained intact and were indistinguishable from control COCs. The cumulus cell area of PCI + GDF9-treated COCs was similar to control ± GDF9 COCs (p>0.99 for all comparisons) (**Fig 7F**). In contrast, the area of PCI-treated COCs was significantly reduced relative to control ± GDF9 COCs and PCI + GDF9-treated COCs (p<0.001 for both comparisons) (**Fig 7F)**. Overall, these data demonstrate that GDF9 fully rescues the loss of matrix integrity secondary to PCSK inhibition and is a likely downstream substrate of PCSK5A in the COC.

## Discussion

Cumulus cells have well-established roles early in ovulation, including support of oocyte maturation and expansion of the COC matrix. However, the key molecules that drive cumulus cell behavior, particularly in later stages leading to follicle rupture, have been less studied. Here, we identified a previously undescribed role of PCSK activity, likely attributable to PSCK5A, in regulating cumulus cell matrix organization and cumulus cell migratory behavior during ovulation. We demonstrated that PCSK inhibition in intact follicles and isolated COCs during the ovulatory process completely disrupted COC matrix integrity, and this was associated with failed follicle rupture. This loss of matrix integrity in COCs occurred during late stages of IVM, indicating a highly temporal role of PCSK activity. Subsequent transcriptomic and proteomic analyses revealed that PCSK inhibition of COCs led to dysregulation in key molecules related to ECM organization, migration, and adhesion pathways consistent with the observed phenotype. Notably, PCSK inhibition was rescued by GDF9 supplementation, suggesting that PCSKs in cumulus cells act via a GDF9-dependent mechanism. The findings lead to a model in which PCSK5A inhibition induces loss of mature GDF9, with subsequent effects on COC matrix remodeling, cumulus cell behavior (adhesion/migration), signaling, and metabolism in the late ovulatory period (**Fig 7G**).

Our study identified novel roles for the PCSKs and their downstream regulators, such as GDF9, in maintaining COC matrix integrity during late ovulation. The PCSKs are broadly expressed across tissue types and cleave various substrates, including transcription factors, receptors, ECM components, and other enzymes^49^. Although previous studies have shown that loss of specific PCSK family members (e.g., *Pcsk3*, *Pcsk6*) causes ovarian dysfunction leading to impaired fertility^50,107^, this study demonstrates that PCSK activity is involved in cumulus cell function. Subsequent findings via RNA *in situ* hybridization and pharmacological inhibition of specific PCSKs indicate that the observed phenotype is primarily driven by PCSK5A, but the exact substrates preferentially cleaved by PCSK5A in cumulus cells remain to be defined. Our multi-omics datasets identified that a suite of pathways dysregulated by PCSK inhibition included TGF-β signaling which is notable given that TGF-β ligands are known PCSK substrates^57–60^. Indeed, the observed disruptions in matrix integrity of PCI-treated COCs could be rescued by supplementation of the oocyte-secreted factor GDF9 which is a member of the TGF-β superfamily and contains a potential site for PCSK-mediated cleavage. During early ovulation, GDF9 is transferred from the oocyte to cumulus cells via transzonal projections to promote initial cumulus cell expansion^108–112^. However, our findings suggest that GDF9 continues to regulate cumulus cell behavior during late ovulation even after retraction of transzonal projections and loss of oocyte-cumulus cell communication. Thus, cumulus cells may also orchestrate their own matrix formation through the processing and activation of GDF9 via PCSK5A. In addition, it is possible that there are other PSCK5A substrates as previous proteolytic analyses have suggested that PCSK5 can cleave key ovulatory proteins, including AMH^113^, BMPs^114^, and metalloproteinases^115,116^. Defining the precise substrates of PSCK5A is an active area of ongoing research.

PCSK inhibition accelerated the normal temporal profile of cumulus cell gene expression during ovulation. Key pathways that were dysregulated by PCSK inhibition included those related to ECM organization, adhesion, and migration, and these were also reflected at the proteome level. Cumulus cell behavior is intricately tied to its surrounding extracellular environment, and altered physical environments may facilitate differential cumulus cell migration^82,117^. During the late stages of ovulation, cumulus cells are known to adopt a more migratory and invasive behavior suggested to promote cumulus cell dispersion^61,95^. Following follicle rupture and release of the COC, the full dispersion of cumulus cells facilitates access for sperm penetration during fertilization^92,118^. These studies align with our ‘omics datasets which demonstrated downregulation of ECM organization and upregulation of migratory pathways with PCSK inhibition. Additionally, we found that PCSK inhibition dysregulated adhesion pathways, leading to a premature reduction in adhesion between cumulus cells compared what is normally observed during late ovulation^61,95^. These findings suggest that PCSK inhibition induces loss of coordination between cumulus cells and their surrounding ECM leading to premature cumulus cell dispersion and subsequent blocking of follicle rupture. Despite all these changes, we observed no effect on oocyte meiotic maturation with PCI treatment demonstrating that PCSK activity is specific to the cumulus cells during late ovulation. Collectively, this study highlights the requirement for synchrony between the oocyte, cumulus cells, and the ECM on a cellular and molecular level for successful ovulation and follicular rupture.

We noted a high correlation between RNA and protein expression in our multi-omic studies with PCI-treated COCs (R=0.898). This is in contrast to other biological contexts where there is low concordance between transcriptomics and proteomics results in cells treated under the same conditions^119–122^, with one study reporting an average correlation of 0.27 between mouse RNA and protein^123^. However, “molecular machines” such as ribosomes and adhesion complexes tend to show high correlation. Consistent with this, translation and adhesion pathways were enriched in both datasets. Intriguingly, we also observed high concordance and enrichment of tumor development pathways (epithelial mesenchymal transition, cell adhesion, and inflammatory response) at the transcript and protein levels. Tumor cells exhibit greater correlation between RNA and protein than non-tumor cells, and genes/proteins with higher correlation are associated with worse outcomes^124^. Ovulation is an inflammation-like process that requires tight orchestration between multiple systems (COC, follicle, ovarian stroma) to occur successfully within a span of hours. Furthermore, cumulus cells acquire migratory and invasive properties in a narrow window just before ovulation, which has been postulated to facilitate follicle wall breakdown and subsequent rupture^61^. Therefore, the high correlation between transcript and protein in the COC during ovulation in this context is reasonable given that the COC must rapidly transcribe RNA and translate protein to accomplish its functions in a highly dynamic setting.

The discovery of the PCSKs and their downstream regulators as mediators of the COC matrix and cumulus cell behavior has significant implications for ovarian function. To our knowledge, this is the first time that PCSK5A has been associated with a reproductive phenotype. Conditional knockouts of *Pcsk5a* in ovarian cell types have not yet been generated and this represents an important avenue of future investigation. However, our work here suggests that genetic polymorphisms of *Pcsk5a* may be associated with anovulatory disorders and infertility. As such, PCSK5A or its regulators could be specifically targeted in the ovary to promote ovulation and continued estrous cycling to benefit reproductive and systemic health. Similarly, downstream regulators of the PCSKs could also be harnessed as non-hormonal contraceptive targets, as PCSK inhibition of follicles does not affect progesterone secretion^47^. In conclusion, this study presents a novel function of the PCSKs to promote sustained matrix integrity and regulate cumulus cell migration via GDF9 during late ovulation.

## Resource Availability

### Lead contact

Requests for further information and resources should be directed to and will be fulfilled by the lead contact, Francesca Duncan.

### Materials availability

This study did not generate new unique reagents.

### Data and code availability

Bulk RNAseq data have been deposited at GEO at GEO: GSE331136. Raw and complete mass spectrometry data sets have been uploaded to the Mass Spectrometry Interactive Virtual Environment (MassIVE) repository, developed by the Center for Computational Mass Spectrometry at the University of California San Diego, and can be downloaded using the following link: https://massive.ucsd.edu/ProteoSAFe/dataset.jsp?task=8bbae4c719834da29605f1fb8aa1842c. (MassIVE ID number: MSV000102085; ProteomeXchange ID: PXD079413). Both the bulk RNAseq and proteomics datasets are publicly available as of the date of publication. This paper also analyzes existing, publicly available data, accessible at GEO at GEO: GSE294534.

All original code has been deposited at Github and is publicly available at https://github.com/RHuang-0/PCSK as of the date of publication. Any additional information required to reanalyze the data reported in this paper is available from the lead contact upon request.

## Acknowledgements

We would like to thank Tim McPhee and Pawat Pattarawat for their assistance with the cumulus cell adhesion and follicle assays, respectively. We also express our gratitude towards Hugh Clarke and Karen Carvalho for helpful discussions. This work was funded by the Gates Foundation (INV-084649, INV-003385, and INV-040475) and the Eunice Kennedy Shriver National Institute of Child Health and Human Development of the National Institutes of Health (T32HD094699). This work was further supported by the Northwestern University NUSeq Core Facility. A.J.D. and C.M.M. acknowledge the support from Arthritis UK (grant: 22277). The content is solely the responsibility of the authors and does not necessarily represent the official views of the National Institutes of Health

## Author contributions

C.E.K., J.P., and F.E.D. conceptualized the study. All authors participated in designing the methodology. C.E.K., R.H., R.M.S., C.D.K., J.Z., and B.S. conducted investigations and formal analyses. C.M.M. and A.J.D. provided resources. C.D.K., B.S., and R.H. were involved in data curation and software. C.E.K., R.H., J.P., and B.S. contributed to data visualization. C.E.K., R.H., J.P., C.D.K., and F.E.D. were involved in writing the original draft of the manuscript. C.E.K., J.P., and F.E.D. reviewed and edited the manuscript. Funding was acquired by C.E.K., R.H., N.P., S.X., D.L.R., B.A.G., and F.E.D. J.P. and F.E.D. supervised this study. F.E.D. was responsible for project administration.

## Supplemental information

Document S1: Sup Figures 1-12, Sup Tables 1 and 2

Sup File 1. Differentially expressed genes between control and PCI-treated COCs at 8h *in vitro* maturation

Sup File 2. Differentially expressed genes between control and PCI-treated COCs at 12h *in vitro* maturation

Sup File 3. Gene ontology

Sup File 4. DIA isolation scheme

Sup File 5. All proteins detected in COCs at 10h *in vitro* maturation by proteomics

Sup File 6. Differentially expressed proteins between control and PCI-treated COCs at 10h *in vitro* maturation

Sup File 7. Matrisome analysis

## Supplemental figure and table legends

**Supplemental Figure 1: PCSK inhibition disrupts COC matrix integrity in a dose dependent manner beginning at 12 h IVM.** Related to Figure 1. Brightfield images of COCs cultured in PCI at increasing concentrations or DMSO control (CTL) for 14-16 h in the EmbryoScope+^TM^ time-lapse incubator.

**Supplemental Figure 2: PCSK inhibition does not affect *in vitro* fertilization outcomes.** Related to Figure 1. Graphs showing the percentage of A) *in vitro* matured CTL, B) *in vitro* matured PCI-treated, and C) *in vivo* matured CTL COCs that reach various stages of embryologic development each day after *in vitro* fertilization. For A-C), data are represented as mean ± SEM and n=42-48 COCs per treatment group. D) Representative immunofluorescent images of blastocysts yielded from CTL or PCI-treated COCs. DNA is shown in blue, alpha-tubulin is shown in green, and rhodamine phalloidin (actin) is shown in red.

**Supplemental Figure 3: PCSK inhibition does not affect the morphokinetics of maturation in mouse oocytes.** Related to Figure 1. A) Representative brightfield images of CTL vs. PCI-treated denuded oocytes across *in vitro* maturation. GV = germinal vesicle. GVBD = germinal vesicle breakdown. PBE = polarbody extrusion. White asterisks denote intact GVs and yellow asterisks denote polarbodies. B) Graphs showing time to germinal vesicle breakdown (GVBD), time to polarbody extrusion (PBE), and overall duration of meiosis in CTL vs. PCI-treated denuded oocytes. Data are represented as mean ± SEM. p > 0.05 for all comparisons by unpaired t-tests. n=24-39 COCs per treatment group. C) Graph showing maturation status of CTL or PCI-treated denuded oocytes after 16h of culture. Data are represented as mean ± SEM. p>0.05 by Welch’s t-test. n=24-39 COCs per treatment group. D) Immunofluorescent images depicting chromosomes (DAPI; blue), spindles (alpha-tubulin; green), and actin (rhodamine phalloidin; red) in CTL or PCI-treated denuded eggs. White asterisks denote spindles and yellow asterisks denote the polar body. Scale bar is 25 µm. Graphs showing E) percent of denuded eggs with bipolar spindles and F) aligned chromosomes after culture with or without PCI. For E) and F), Data are represented as mean ± SEM, p>0.05 by Fisher’s exact test, and n=3 biological replicates (24-39 pooled COCs per treatment group).

**Supplemental Figure 4: *Pcsk3* and *Pcsk5a* transcripts are similarly expressed in cumulus cells.** Related to Figure 2. A) 20x scans of whole mouse ovaries fixed and stained for *Pcsk3*, *5a*, *5b*, or *6* using RNA *in situ* hybridization (RNAscope). 20x scans of RNAscope positive and negative controls (*Ppib* and *Dapb*, respectively) also shown. B) Representative deconvoluted images from RNA *in situ* hybridization depicting expression of *Pcsk3*, *5a*, *5b*, and *6* mRNA transcripts (red dots) in COCs at 0 h, 4 h, 8 h, or 11 h post-hCG injection. C) Quantification of *Pcsk3*, *5a*, *5b*, or *6* expression shown separately by gene. Data are represented as mean ± SEM. p-values as follows: * < 0.05, ** < 0.01, *** < 0.001, **** < 0.0001 by one-way ANOVA. n=3-5 biological replicates per *Pcsk*.

**Supplemental Figure 5: PCSK5 is highly abundant in cumulus cells.** Related to Figure 2. A) 20x scans of whole mouse ovaries fixed and stained for PCSK3 and PCSK5 using immunohistochemistry. Scale bar = 200 µm. 20x scans of non-immune control sections for each antibody also shown. B) Representative deconvoluted images from immunohistochemistry depicting abundance of PCSK3 and PCSK5 protein in COCs at 0 h, 4 h, 8 h, or 11 h post-hCG injection.

**Supplemental Figure 6: PCI (pan-PCSK inhibitor) has a more severe effect on matrix integrity than FII (PCSK3 inhibitor).** Related to Figure 2. A) Brightfield images of COCs cultured in PCI- or FII-treated media or DMSO control (CTL) for up to 16 h in the EmbryoScope+^TM^ time-lapse incubator. B) Immunofluorescent images depicting chromosomes (DAPI; blue), spindles (alpha-tubulin; green), and actin (rhodamine phalloidin; red) in CTL, FII-, or PCI-treated COCs after denuding of cumulus cells. White asterisks denote spindles and yellow asterisks denote the polar body. Scale bar is 25 µm. C) Graph showing maturation status of FII- and PCI-treated COCs relative to CTL COCs. Data are represented as mean ± SEM. p>0.05 by Fisher’s exact test. n=11-26 biological replicates.

**Supplemental Figure 7: Variability of transcriptomic and proteomic samples.** Related to Figures 3 and 4. A) PCA plot of CTL vs. PCI-treated COCs across a time course of *in vitro* maturation. B) PLS-DA of CTL vs. PCI-treated COCs at 10 h after onset of *in vitro* maturation.

**Supplemental Figure 8: Pathways up- and down-regulated with PCSK inhibition in 12 h COCs are similar to those of 8 h PCI-treated COCs.** Related to Figure 3. A) Dot plot showing pathways downregulated with PCI treatment at 12 h. B) Dot plot showing pathways upregulated with PCI treatment at 12 h. C) Heatmap of top genes associated with metabolism that are upregulated by PCI treatment. D) Dot plot showing pathways common between 8 h PCI and 12 h CTL COCs.

**Supplemental Figure 9: Pathways related to translation, metabolism, immune function, and surveillance are enriched in control COCs at 10h *in vitro* maturation.** Related to Figure 4. Heatmap showing expression of all proteins dysregulated with PCI treatment in CTL vs. PCI-treated COCs.

**Supplemental Figure 10: Corresponding gene expression of proteins dysregulated by PCI treatment from bulk RNAseq of control vs. PCI-treated COCs.** Related to Figures 5-7. A) Box plots of *Tsg-6*, *Plau,* and *Plaur* across IVM in control (CTL) and PCI-treated COCs. B) Box plots of adhesion-related genes *Itga5* and *Sema7a* across IVM in CTL and PCI-treated COCs. C) Box plots of TGF-β signaling related genes *Tgfbr1*, *Fst*, *Dab2*, and *Bmpr2* across IVM in CTL and PCI-treated COCs. For all graphs, data are represented as mean ± SD. * p < 0.05 for all graphs by Wilcoxon signed-rank test. n=3 biological replicates of 10 pooled COCs per treatment and timepoint.

**Supplemental Figure 11: Total abundance of hyaluronan staining is unchanged with PCI treatment.** Related to Figure 5. Bar graph showing the total area of hyaluronan stained with HABP normalized by the total cumulus cell area. Data are represented as mean ± SEM. * p < 0.05 by two-way ANOVA. n=13-21 biological replicates.

**Supplemental figure 12: Cumulus cell adhesion is dysregulated with PCI treatment.** Related to Figure 6. Violin plots showing abundance of adhesion-related proteins A) IGFBP5 and B) PTK2B in CTL vs. PCI-treated COCs at 10 h IVM. For A) and B), data are represented as mean ± SEM. ** q < 0.01 by t-test with Storey’s method. n=4 biological replicates, where each replicate has 150 COCs per treatment group. C) Schematic showing predicted behavior of cumulus cells in a weak PCI-treated ECM. D) Relative adhesion of CTL vs. PCI-treated COCs on fibronectin-coated plates. Data are represented as mean ± SEM. * p < 0.05 by unpaired t-test. n=8 biological replicates per treatment group.

**Supplemental Table 1:** Genes driving pathways related to cell adhesion, cell movement, or metabolism which are differentially expressed with PCI treatment (mined from Enrichr).

**Supplemental Table 2:** Oocyte markers are stable with PCI treatment.

## Methodology

### Experimental model

#### Mice

Prepubertal and adult (6-12 weeks old) female CD-1 mice (Inotiv, West Lafayette, IN, USA) were housed in a controlled barrier facility at the Northwestern University Center for Comparative Medicine (Chicago, IL, USA) or Rutgers University (Piscataway, NJ, USA). These facilities were kept at constant temperature and humidity with a 14h light/10h dark cycle. Up to five mice were housed per cage. Food and water were provided ad libitum. Mice received chow excluding soybean and alfalfa meal to reduce exposure to phytoestrogens (Teklad Global 2916 chow, Inotiv, West Lafayette, IN, USA). Following arrival at these facilities, mice were acclimated for at least one week prior to use. All experiments were approved by the Institutional Animal Care and Use Committee at Northwestern University and Rutgers University. This study was also performed in accordance with the National Institute of Health Guide for the Care and Use of Laboratory Animals.

### Method details

#### Encapsulated in vitro follicle growth and ex vivo ovulation

Ovaries were harvested from prepubertal female CD-1 mice aged 16 days old. Insulin gauge needles (BD Biosciences, Franklin Lakes, NJ, USA) were utilized to puncture the ovaries and release multi-layered secondary follicles. Follicles were collected in Leibovitz’s 1x (L15) media (Thermo Fisher Scientific, Waltham, MA, USA, #11415-144) containing 1% (v/v) fetal bovine serum (FBS; Peak Serum Inc., Wellington, CO, USA). Intact multi-layered secondary follicles were placed individually into 5 µL alginate hydrogel beads. To crosslink the alginate polymers, beads were placed in a 50 mM CaCl_2_ (Thermo Fisher Scientific) and 140 mM NaCl (Thermo Fisher Scientific) solution for 2 minutes. Crosslinked beads were then cultured in 96 well-plates. Each well housed one bead and 100 µL growth media containing 50% (v/v) α-MEM (1x) + GlutaMAX™ media (α-MEM; Thermo Fisher Scientific, #32561037) and 50% (v/v) Ham’s F-12 Nutrient Mix, GlutaMAX™ Supplement F-12 Glutamax media (F12; Thermo Fisher Scientific, #31765035) plus 3 mg/mL bovine serum albumin (BSA; Sigma-Aldrich, St. Louis, MO), 10 mIU/mL recombinant follicle stimulating hormone (rFSH; gift from Organon, Jersey City, NJ), 2 mg/mL bovine fetuin (Sigma-Aldrich, #F3385), and 5 µg/mL insulin-transferrin-selenium (Sigma-Aldrich, #I1884-1VL). Follicles were cultured in humified conditions at 37°C and 5% CO_2_, with 50 µL growth media replaced every other day.

After 8 days of culture, antral follicles sized 300-350 µm in diameter were selected for *ex vivo* ovulation. These follicles were removed from the alginate beads in L15 media supplemented with 1% FBS and 10 IU/mL alginate lyase extracted from *Flavobacterium multivorum* (Sigma-Aldrich, #A1603). Follicles were then cultured in maturation media containing 50% (v/v) α-MEM and 50% (v/v) F-12 with 10% FBS, 1.5 IU/mL human chorionic gonadotropin (hCG; Sigma-Aldrich), 10 ng/mL epidermal growth factor (EGF; BD Biosciences) and 10 mIU/mL rFSH for 14 h in PCI or DMSO control.

#### PCSK inhibitor treatment

PCSK inhibitors utilized in this study include proprotein convertase inhibitor (PCI; Sigma-Aldrich, #537076) and Furin Inhibitor II (FII; Sigma-Aldrich, #SCP0148). For all experiments testing inhibitor treatment, *ex vivo* ovulation of follicles or *in vitro* maturation of COCs was performed in the presence of 10 µM PCSK inhibitor prepared from a 20 µM stock solution in dimethyl sulfoxide (DMSO; Sigma-Aldrich, #M1404). Inhibitors were added at the beginning of the culture period and remained in the media through the duration of culture. DMSO at a concentration of 1/2000 was used as a negative control. In rescue experiments with GDF9, recombinant murine GDF9 was added at a concentration of 500 ng/mL (R&D Systems, Minneapolis, MN, USA, #739-G9-010).

#### RNA extraction and RT-qPCR

Antral follicles in maturation media containing DMSO control or 10 µM PCI were collected at 0 h, 4 h, or 8 h after exposure to hCG. The PicoPure RNA Isolation Kit (Thermo Fisher Scientific, #KIT0204 or KIT0214) was utilized to extract RNA from follicles. The Superscript III First-Strand Synthesis System with random hexamer primers (Invitrogen) was used to reverse transcribe extracted RNA into cDNA. cDNA was then stored at -80°C until RT-qPCR was performed.

cDNA was plated into a 96-well plate with the Power SYBR Green PCR Master Mix (Thermo Fisher Scientific). The primer sequences (5’ for 3’) for *Has2* were as follows:

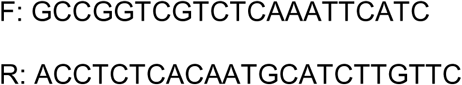

RT-qPCR was conducted using the StepOnePlus Real-Time PCR system (Thermo Fisher Scientific). The RT-qPCR thermocycler ran as follows: 10 min at 95°C; 15 seconds at 95°C and 40 seconds at 60°C x 40 cycles; and melting curve stage for evaluation of primer specificity. The relative expression of each gene was calculated using the following equation:

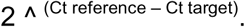

#### Ovarian hyperstimulation and cumulus-oocyte-complex (COC) collection

Mice received an intraperitoneal (IP) injection of 5 IU pregnant mare serum gonadotropin (PMSG; ProSpec-Tany TechnoGene, East Brunswick, NJ, #HOR-272) to recruit follicles. 44-46 h post PMSG injection, ovaries were dissected and placed in pre-warmed L15 medium (Thermo Fisher Scientific, #11415-144) containing 3 mg/mL polyvinylpyrrolidone (PVP; Sigma-Aldrich, #P2307) and 0.5% (v/v) penicillin-streptomycin (PS; Life Technologies Corporation, Grand Island, NY, #15140122) (L15/PVP/PS). Ovaries were then poked with an insulin syringe needle (Fisher Scientific, Hampton, NH, #26027) to release intact COCs from antral follicles. Collected COCs were transferred to a dish containing L15/PVP/PS media containing 2.5 µM milrinone (Sigma-Aldrich, #M4659) to prevent spontaneous resumption of meiosis. For culture of denuded oocytes, COCs were mechanically denuded to remove surrounding cumulus cells. Denuded oocytes were allowed to recover in α-MEM (1x) + Glutamax media (α-MEM) (Thermo Fisher Scientific, #32561037) containing PS and bovine serum albumin (BSA; Sigma-Aldrich, #A3311) (α-MEM/PS/BSA) at 37°C and humidified atmosphere of 5% CO_2_ for up to 3 h prior to incubation. To reduce animal-specific variability, denuded oocytes or COCs were pooled before being distributed to culture dishes.

### In vitro maturation (IVM) of COCs

#### EmbryoScope+^TM^ time-lapse incubator

For culture of denuded oocytes, α-MEM/PS/BSA was utilized as the culture medium. For culture of intact COCs, media specifically formulated to promote COC expansion was utilized: α-MEM containing 5% (v/v) fetal bovine serum (FBS; Thermo Fisher Scientific, #10082139), 20 mM HEPES buffer (Sigma-Aldrich, #H6147), 0.25 mM pyruvate (Pyr; Sigma-Aldrich, #P2256), and 0.02% (w/v) mouse epidermal growth factor (mEGF; Sigma-Aldrich, #SRP3196) (α-MEM/FBS/HEPES/Pyr/mEGF). The day prior to COC collection, the appropriate culture medium was loaded into microwells of the EmbryoSlide^TM^ culture dishes. These dishes were overlayed with 1.6 mL mineral oil and equilibrated overnight (9-24 h) in the EmbryoScope+^TM^. Just before incubation, denuded oocytes or COCs were washed in four large drops of equilibrated α-MEM/PS/BSA or a-MEM/FBS/HEPES/Pyr/mEGF media, respectively, to remove residual milrinone and allow meiotic resumption. One denuded oocyte or COC was loaded in each microwell. Denuded oocytes and COCs were cultured for 16 h in the EmbryoScope+^TM^ in humified conditions at 37°C and 5% CO_2_. COCs were transferred to pre-warmed L15/PVP/PS media and denuded at the completion of IVM to check maturation status. The EmbryoScope+^TM^ captured images of each well within the EmbryoSlide^TM^ dish at 10-min intervals in 11 different focal planes. All images were taken using low-intensity red LED illumination with less than 0.5 s of light exposure per image, mimicking conditions used for human embryos in ART.

After 14-16 h culture in the EmbryoScope+^TM^, COCs were removed from individual Embryoslide^TM^ wells and transferred to a separate petri dish. Transferred COCs were imaged using the EVOS FL Auto imaging system. COC area calculated using Fiji. Individual COCs were outlined using the freehand selection tool, and then the total area of each selection was measured.

#### Standard incubator

Center well dishes (Corning, Corning, NY, #353037) were loaded with 1 mL α-MEM/PS/BSA or α-MEM/FBS/HEPES/Pyr/mEGF media for culture of denuded oocytes or COCs, respectively. Media was equilibrated for at least 3 h prior to culture. Denuded oocytes and COCs were washed free of milrinone as described above before being loaded into center well dishes and cultured for 16 h in humified conditions at 37°C and 5% CO_2_. COCs were transferred to pre-warmed L15/PVP/PS media and denuded at the completion of IVM to check maturation status.

### Viability/cytotoxicity assay

COC collection was performed as described above. DMSO control and PCI-treated COCs were added to separate 35 mm MatTek dishes with No. 1 uncoated coverslip (Fisher Scientific, #NC0179740) each containing a single 85 µL drop of α-MEM/FBS/HEPES/Pyr/mEGF under mineral oil, based on assigned treatment group. Collected COCs were cultured in the standard incubator as described above. Following culture, the Live/dead™ Viability/Cytotoxicity Kit, for mammalian cells (Invitrogen, #L3224) was used to assess viability of cumulus cells. 4 mM calcein AM (live cell dye) and 2 mM ethidium homodimer (dead cell dye) were added to the media drops containing COCs to final concentrations of 1 µM and 2 µM, respectively. COCs were then incubated for an additional 30 min in the standard incubator before imaging on the Leica SP5 confocal. Two independent trials were performed.

### Immunofluorescence

#### Staining

Denuded gametes (including COCs denuded at the conclusion of IVM) that extruded polar bodies were fixed at 37°C for 20 min in 3.8% (v/v) paraformaldehyde (PFA; Electron Microscopy Sciences, Hatfield, PA, #157-4-100) supplemented with 0.1% TritonX-100 (TX-100; Alfa Aesar, Haverhill, MA, #A16046) to improve spindle visualization. Following fixation, gametes were washed 2 x 5 min in blocking buffer containing 1X PBS (Thermo Fisher Scientific, #J61196-AP), 0.01% (v/v) Tween-20 (Sigma-Aldrich, #P1379), 0.02% (w/v) sodium azide (NaN_3_; Sigma-Aldrich, #71289), and 0.3% (w/v) BSA. Gametes were then permeabilized for 15 min at room temperature in 1x PBS/0.1% TX-100/0.02% NaN_3_/0.3% BSA Following permeabilization, gametes were washed 2 x 5 min in blocking buffer. Gametes were then incubated for 2 h at room temperature with rhodamine phalloidin (1:50; Invitrogen, #R415) and α-tubulin (11H10) rabbit mAb 488 (1:100; Cell Signaling Technology, Danvers, MA, #5063S) to visualize actin and microtubules, respectively. After staining, gametes were washed 3 x 20 min in blocking buffer. Gametes were mounted in Vectashield Plus Antifade Mounting Medium with DAPI (4,6-diamidino-2-phenylindole; Vector Laboratories, Burlingame, CA, #H-2000) on 1.0 mm thick microscope slides. Two independent trials each were performed for PCI vs. FII-treated COCs. Three independent trials each were performed for COCs and denuded oocytes ± PCI treatment.

#### Confocal imaging and image processing/analysis

Images were captured on the Lecia SP5 inverted laser scanning confocal microscope (Leica Microsystems, Wetzlar, Germany) at 40x magnification with the 405, 488, and 542 nm lasers. Z-stack thickness was 1 µm. LAS AF (Leica Microsystems) and FIJI (National Institutes of Health, Bethesda, MD) were used for image processing and analysis, respectively.

For evaluation of morphology where protein expression was not directly compared between images, the gain, laser power, and brightness were adjusted between images for best visualization of the chromosomes and cytoskeleton. Spindle polarity was determined by counting the number of spindle poles within each egg. A spindle pole number of 2 was considered normal (bipolar). Chromosome alignment was determined by evaluating the localization of chromosomes relative to each other and the spindle. Chromosomes which did not contact other chromosomes within the metaphase plate were considered misaligned.

### In vitro fertilization (IVF) and embryo culture

COCs were cultured in a standard incubator with or without PCI for 12 h as described above. After 11.5 h of IVN COC culture, spermatozoa were collected from the caudal epididymides of male CD1 mice (Inotiv) aged 8-10 weeks old. Collected sperm were incubated in HTF media (100 mM NaCl, 4.7 µm KCl, 200 mM MgS0_4_, 400 µm KH_2_PO_4_, 5 nM CaCl_2_, 2.77 mM glucose, 16 µm sodium lactate, 336 µm sodium pyruvate, 200 µm penicillin G, 70 µM streptomycin, 25mM NaHCO_3_, 0.01% (w/v) phenol red) supplemented with 4 mg/mL BSA in an IVF center well dish overlayed with embryo safe mineral oil for 30 min at 37°C and 5% CO_2_ to permit dispersion. During this 30 min sperm incubation period, *in vivo* matured COCs were collected from the ampulla of female mice hyperstimulated with hCG 46 h prior.

At the completion of 12h IVM, *in vitro* matured COCs were washed in HTF+BSA and added to an IVF center well dish containing fresh HTF+BSA. Motile sperm were added to center well dishes containing *in vitro* control, *in vitro* PCI, or *in vivo* control COCs. The combined COCs and sperm were incubated for 6 h at 37°C and 5% CO_2_ to allow fertilization. After the 6 h incubation period, COCs were washed in Potassium-supplemented Simplex Optimized Medium (KSOM) to remove any attached sperm and transferred to EmbryoScope+^TM^ dishes containing KSOM for embryo development. After 120 h of culture, blastocysts were fixed for 1 h at room temperature in 3.8% PFA + 0.1% Tx-100 + 0.03% (w/v) PVP. The remainder of the fixation was performed as described above until mounting. Just before mounting in Vectashield Plus Antifade Mounting Medium with DAPI (4,6-diamidino-2-phenylindole; Vector Laboratories, #H-2000), blastocysts were washed in increasing concentrations of Vectashield diluted in water (25%, 50%, 75%, 90% (v/v)) to prevent collapse. Blastocysts were stained for alpha-tubulin and F-actin as described above. Two independent trials were performed.

### Analysis of meiotic progression

At the conclusion of IVM, COCs were denuded using 0.25 mg/mL testicular hyaluronidase (Sigma-Aldrich, #H4272) prior to scoring. The meiotic stage of all gametes was then scored based on the following morphological criteria. Oocytes that arrested at prophase I of meiosis, distinguished by the presence of an intact nuclear envelope, were designated as germinal vesicles (GV). Oocytes that underwent GVBD but did not extrude a polar body were either arrested at pro-metaphase I or metaphase I (MI) and classified as GVBD/MI. Progression to MI and was confirmed by alignment of chromosomes at the metaphase plate without a polar body (PB) by immunofluorescence. Gametes which extruded PBs were designated as eggs arrested at metaphase II (MII).

Morphokinetics of meiotic maturation were measured as previously described^53,125,126^. The time-lapse images captured by the EmbryoScope+^TM^ were utilized to determine time to the key meiotic events described below for denuded oocytes. In COCs, cumulus cells obstructed the view of the oocyte and precluded measurement of these parameters. The time at which COCs or denuded oocytes were placed in the EmbryoScope+^TM^ for IVM and imaging was designated as 0 h. Time to GVBD was marked by complete breakdown of the germinal vesicle and disappearance of the nuclear envelope. Time to first PB extrusion was designated as the point at which cytokinesis was completed and there were clear borders between the plasma membranes of the oocytes and play body. The duration of meiosis I was calculated as the difference in time between GVBD and PB extrusion. Three independent trials were performed.

### RNA in situ hybridization

The RNAscope 2.5 HD Red Assay (Advanced Cell Diagnostics (ACD), Newark, CA, USA, #322360) was used to detect the expression of *Pcsk3* (ACD, #864041), *Pcsk5a* (ACD, #1241481-C1), *Pcsk5b* (ACD, #1241471-C1), and *Pcsk6* (ACD, #593361) mRNA in whole formalin-fixed-paraffin-embedded (FFPE) ovary sections. *Ppib* (ACD, #313911) and *Dapb* (ACD, #310043) were used as positive and negative control probes, respectively. Ovary sections were incubated in target or control probes for 2 h at 40°C followed by counterstaining with 50% Gill’s hematoxylin (Fisher Scientific, Hampton, NH, USA, #22-021-416) and ammonia water. The ACD website contains the stepwise protocol for this assay (https://acdbio.com/sites/default/files/322360-USM%20RNAscope%202.5%20HD%20RED%20Pt2_11052015.pdf).

Whole ovary scans (20x magnification) and images highlighting COCs (40x magnification) were captured using the EVOS FL Auto imaging system, then color deconvoluted using Fiji for improved visualization and quantification. To determine the RNA signal within cumulus cells, regions of interest (whole COC and the oocyte alone) were outlined using the freehand selection tool. The area of the whole COC and oocyte alone were determined by measuring the thresholded area of DAPI signal within the outlined regions of interest. Next, the amount of RNA within each region of interest was determined by measuring the thresholded area of transcript signal. Thresholds for DAPI and RNA transcripts were kept constant for all images within a trial. The area of the cumulus cell compartment was calculated by subtracting the oocyte DAPI area from the whole COC DAPI area. The amount of RNA within the cumulus cell compartment was calculated by subtracting the oocyte transcript area from the whole COC transcript area. The amount of RNA within cumulus cells was then normalized by the total area of the cumulus cell compartment. Three independent trials were performed.

### Immunohistochemistry

FFPE ovary sections were deparaffinized in Citrisolv and rehydrated in decreasing concentrations of ethanol. 1X Reveal Decloaker (Biocare Medical, Concord, CA, USA, #RV1000) was used for head-induced antigen retrieval. Briefly, slides containing FFPE ovary sections were microwaved at 50% power for 2 min and 10% power for 10 min in the Reveal Decloaker. Following antigen retrieval, slides were incubated in 3% (v/v) hydrogen peroxide for 15 min to block endogenous peroxidase activity. An avidin/biotin blocking kit (15 min in each blocking solution) was utilized to block endogenous avidin and biotin. Slides were then incubated for 1 h in 10% (v/v) normal goat serum and 3% (w/v) BSA in Tris-buffered saline to block non-specific antigens. Each blocking step was performed at room temperature in a humidified chamber. Following goat serum block, slides were incubated with rabbit anti-mouse PCSK3 primary antibody (1:1000; Thermo Fisher Scientific, #PA1-062) or PCSK5 primary antibody (1:10000; Proteintech, Rosemont, IL, #16470-1-AP) diluted in 3% BSA overnight at 4°C in a humidified chamber. After washing in TBS with 3% BSA, slides were incubated with biotinylated anti-rabbit IgG secondary antibody (1:200; Vector Laboratories, Newark, CA, # PK- 6101) diluted in 3% (w/v) BSA for 1 h at room temperature in a humidified chamber. The Vectastain Elite ABC Kit (Vector Laboratories, Newark, CA, #PK-6101) was used to amplify the signal before detection and the DAB Peroxidase (HRP) Substrate Kit was used to detect the signal with 3,3’-diaminobenzidine 25 (Vector Laboratories, Newark, CA, #SK-4100). After signal detection with DAB, slides were counterstained with Harris hematoxylin (EK Industries, Joliet, IL, #4797). Increasing concentrations of ethanol and Citrisolv were used to dehydrate and clear the tissue, respectively. Sections were mounted with Cytoseal XYL.

Whole ovary scans (20x magnification) and images highlighting COCs (40x magnification) were captured using the EVOS FL Auto imaging system, then color deconvoluted using Fiji for improved visualization and quantification. To determine the protein signal within cumulus cells, regions of interest (whole COC and the oocyte alone) were outlined using the freehand selection tool. The area of the whole COC and oocyte alone were determined by measuring the thresholded area of hematoxylin signal within the outlined regions of interest. Next, the amount of protein within each region of interest was determined by measuring the thresholded area of DAB signal. Thresholds for hematoxylin and DAB transcripts were kept constant for all images within a trial. The area of the cumulus cell compartment was calculated by subtracting the oocyte hematoxylin area from the whole COC hematoxylin area. The amount of protein within the cumulus cell compartment was calculated by subtracting the oocyte protein signal area from the whole COC protein signal area. The amount of protein within cumulus cells was then normalized by the total area of the cumulus cell compartment. Two independent trials were performed.

### Bulk RNA-sequencing

#### Sample preparation

COCs were collected from mouse ovaries and cultured in α-MEM/FBS/HEPES/Pyr/mEGF with DMSO or 10 µM PCI as described above. COCs were cultured in the standard incubator for 0 h, 4 h, 8 h, or 12 h (10 COCs per group). RNA extraction was performed using the RNeasy Micro Kit (Qiagen, Germantown, MD, #74004). At the end of the designated culture period, COCs were washed in 1x PBS before transferring to 1% (v/v) beta-mercaptoethanol in Buffer RLT. COCs were then snap frozen and stored at -80°C before extraction. Three independent time courses were performed. Extraction was performed for all samples simultaneously.

#### General mRNA library construction

To generate RNA-sequencing libraries, RNA quality was determined with the Agilent Bioanalyzer 2100, accepting RNA integrity numbers (RIN) of > 7, and quantified using Qubit. Directional mRNA libraries were prepared using Illumina TruSeq mRNA Sample Preparation Kits per manufacturer’s instructions. Briefly, polyadenylated mRNAs were captured from total RNA using oligo-dT selection. Next, samples were converted to cDNA by reverse transcription, and each sample was ligated to Illumina sequencing adapters containing unique barcode sequences. Barcoded samples were then amplified by PCR and the resulting cDNA libraries were quantified using qPCR. Finally, equimolar concentrations of each cDNA library were pooled and sequenced on the Illumina HiSeq 4000.

#### RNA-seq and data pre-processing

The stranded mRNA-seq was conducted in the Northwestern University NUSeq Core Facility. Briefly, total RNA samples were checked for quality using RINs generated from Agilent Bioanalyzer 2100. RNA quantity was determined with Qubit fluorometer. The Illumina TruSeq Stranded mRNA Library Preparation Kit was used to prepare sequencing libraries of high-quality RNA samples (RIN>7). The Kit procedure was performed without modifications. This procedure includes mRNA enrichment and fragmentation, cDNA synthesis, 3’ end adenylation, Illumina adapter ligation, library PCR amplification and validation. Illumina NovaSeq X NGS Sequencer was used to sequence the libraries with the production of paired-end 150 bp reads.

The quality of reads, in FASTQ format, was evaluated using FastQC. Reads were trimmed to remove Illumina adapters from the 3’ ends using cutadapt 26. Trimmed reads were aligned to the Mus musculus genome (mm39) using STAR^127^. Read counts for each gene were calculated using htseq-count^128^ in conjunction with a gene annotation file for mm39 obtained from Ensembl (http://useast.ensembl.org/index.html). Gene-level count matrices were imported into R for downstream analysis. Only protein-coding genes were retained, and genes lacking annotation were removed. To avoid duplicate gene entries, duplicate gene symbols were collapsed, retaining unique gene names.

#### Data analysis

Principal component analysis (PCA) was performed in R using the PCAtools^129^. Samples were annotated by time point and treatment condition. For visualization, treatment groups were defined as baseline (0 h), 4 h control, 4 h PCI, 8 h control, 8 h PCI, 12 h control, and 12 h PCI. PCA biplots were generated with samples colored by the experimental group, and ellipses were added to visualize clustering patterns across replicates and conditions. Heatmaps were generated in R using the ComplexHeatmap^130,131^ package to visualize expression patterns of selected genes of interest across samples. Expression values were standardized by row using Z-score scaling to emphasize relative expression changes across samples. Heatmaps were generated using hierarchical clustering of genes and samples.

To evaluate overall similarity among samples, pairwise sample correlation matrices were computed in R. In one analysis, Pearson correlation coefficients were calculated across the full expression matrix and visualized using pheatmap^132^, with sample annotations corresponding to time point and treatment status. In a second analysis, the top 1,000 most variable genes, ranked by variance across samples, were selected to reduce noise and improve the separation of the biological signal. Pearson correlation coefficients were then calculated among samples using this reduced matrix and visualized as a clustered heatmap.

Differential expression analysis was performed in R using the DESeq2 package^133^. For each time point, differential expression was tested between the PCI and control groups using a design formula of ∼ Ctrl. Genes were ranked by adjusted p-value, and significantly differentially expressed genes were defined as those with adjusted p-value <0.05. Full differential expression tables and significant gene lists were exported for downstream analyses.

Gene ontology analysis was performed using Enrichr^134–136^. All upregulated genes for each comparison of interest (e.g. 8 h control COCs vs. 8 h PCI-treated COCs) were entered into the Enrichr software. The gene ontology results from three Enrichr libraries were utilized for pathway analysis: GO Biological Process, Reactome Pathways 2024, and KEGG 2021 Human. Only pathways with adjusted p-values <0.05 were included in the analysis. Enriched pathways were sorted by descending adjusted p-value, and the top ten pathways from each library were considered. These pathways were then categorized by general biological function. This process was repeated for all statistically significant downregulated genes from each comparison.

### Proteomics

#### Sample preparation

COCs were collected from 60 mice hyperstimulated with 5 IU PMSG. COCs from 20 mice were pooled, then split into two groups of 150 COCs, which yielded three biological replicates of control and PCI-treated COCs. Extra COCs from each group of 20 mice were pooled for a fourth biological replicate. COCs were then cultured in α-MEM/FBS/HEPES/Pyr/mEGF with DMSO or 10 µM PCI in the standard incubator as described above for 10 h. After culture, COCs were washed in four large drops of Dulbecco’s Phosphate Buffered Saline without calcium or magnesium (DPBS-/-, Thermo Fisher Scientific, #14-190-250) supplemented with 0.4% PVP (Sigma-Aldrich, #P2307). COCs were then snap frozen in minimal volume on dry ice and stored at -80°C until further processing. Four biological replicates each of control and PCI-treated COCs were processed and subjected to mass spectrometric analysis.

#### Protein digestion and desalting

Samples (∼10 µL) were brought to the same overall volume of 50 µL with water. All samples were reduced using 20 mM dithiothreitol in 50 mM TEAB at 50°C for 10 min, cooled to room temperature (RT) and held at RT for 10 min, and alkylated using 40 mM iodoacetamide in 50 mM TEAB at RT in the dark for 30 min. Samples were acidified with 12% (v/v) phosphoric acid to obtain a final concentration of 1.2% phosphoric acid. S-Trap buffer consisting of 90% (v/v) methanol in 100 mM TEAB at pH ∼7.1, was added and samples were loaded onto S-Trap mini spin columns (Protifi). The entire sample volume was spun through the S-Trap mini spin column at 4,000 x g and RT, binding the proteins to the micro spin columns. Subsequently, S-Trap micro spin columns were washed twice with S-Trap buffer at 4,000 x g and RT and placed into clean elution tubes. Samples were incubated for one-hour at 47°C with sequencing grade trypsin (Promega, San Luis Obispo, CA) dissolved in 50 mM TEAB at a 1:25 (w/w) enzyme:protein ratio. An additional aliquot of trypsin dissolved in 50 mM TEAB was added and samples were digested overnight at 37°C.

Peptides were sequentially eluted from S-Trap micro spin columns with 50 mM TEAB, 0.5% (v/v) formic acid (FA) in water, and 50% (v/v) acetonitrile (ACN) in 0.5% FA. After centrifugal evaporation, samples were resuspended in 0.2% FA in water and desalted with in-house packed C_18_ Stage-tips. The desalted elutions were then subjected to an additional round of centrifugal evaporation and re-suspended in 20 µg of 0.1% FA in water. Two microliters of each sample was diluted with 2% ACN in 0.1% FA to obtain a dilution of 1:10. One microliter of indexed Retention Time Standard (iRT, Biognosys, Schlieren, Switzerland) was added to each sample, thus bringing up the total volume to 20 µL^137^.

#### Mass spectrometric analysis

Reverse-phase HPLC-MS/MS analyses were performed on a Dionex UltiMate 3000 system coupled online to an Orbitrap Exploris 480 mass spectrometer (Thermo Fisher Scientific, Bremen, Germany). The solvent system consisted of 2% ACN, 0.1% FA in water (solvent A) and 80% ACN, 0.1% FA in ACN (solvent B). Digested peptides (1:10 dilution) were loaded onto an Acclaim PepMap 100 C_18_ trap column (0.1 x 20 mm, 5 µm particle size; Thermo Fisher Scientific) over 5 min at 5 µL/min with 100% solvent A. Peptides were eluted on an Acclaim PepMap 100 C_18_ analytical column (75 µm x 50 cm, 3 µm particle size; Thermo Fisher Scientific) at 300 nL/min using the following gradient: linear from 2.5% to 24.5% (v/v) of solvent B (in solvent A) in 125 min, linear from 24.5% to 39.2% of solvent B (in solvent A) in 40 min, up to 98% of solvent B (in solvent A) in 1 min, and back to 2.5% of solvent B (in solvent A) in 1 min. The column was re-equilibrated for 30 min with 2.5% of solvent B (in solvent A), and the total gradient length was 210 min. Each sample was acquired in data-independent acquisition (DIA) mode^138–140^. Full MS spectra were collected at 120,000 resolution (Automatic Gain Control (AGC) target: 3 x 10^6^ ions, maximum injection time: 60 ms, 350-1,650 *m/z*), and MS2 spectra at 30,000 resolution (AGC target: 3 x 10^6^ ions, maximum injection time: Auto, Normalized Collision Energy (NCE): 30, fixed first mass 200 *m/z*). The isolation scheme consisted of 26 variable windows covering the 350-1,650 *m/z* range with an overlap of 1 *m/z* (Sup File 4)^140^.

#### DIA-MS data processing and statistical analysis

DIA data were processed in Spectronaut v20 (version 20.4.260109.92449) using directDIA. Data extraction parameters were set as dynamic and non-linear iRT calibration with precision iRT was selected. Data were searched against the *Mus musculus* reference proteome with 54,739 entries (UniProtKB-SwissProt), accessed on 03/04/2025. Trypsin/P was set as the digestion enzyme and two missed cleavages were allowed. Cysteine carbamidomethylation was set as a fixed modification while methionine oxidation and protein N-terminus acetylation were set as dynamic modifications. Identification was performed using 1% precursor and protein q-value. Quantification was based on the peak areas of extracted ion chromatograms (XICs) of 3 – 6 MS2 fragment ions, specifically b- and y-ions, with q-value sparse data filtering, iRT profiling (**Sup File 5**), and local normalization applied. Differential protein expression analysis comparing PCI vs control was performed using a paired t-test, and p-values were corrected for multiple testing, using the Storey method^141^. Specifically, groupwise testing corrections were applied to obtain q-values. For secretome analysis, protein groups with at least two unique peptides, q-value <0.05, and absolute Log_2_(fold-change) >0.58 are significantly altered (**Sup File 6**).

For ontology of proteins detected in control COCs alone, the top 150 proteins in control COCs were entered into the Enrichr software. The gene ontology results from three Enrichr libraries were utilized for pathway analysis: GO Biological Process, Reactome Pathways 2024, and KEGG 2021 Human. Only pathways with adjusted p-values <0.05 were included in the analysis. Enriched pathways sorted by descending adjusted p-value, and the top ten pathways from each library were considered. These pathways were then manually categorized by general biological function. This process was repeated for all statistically significant downregulated genes from each comparison. For ontology of differentially expressed proteins, proteins were manually categorized directly based on general biological function.

### Scaffold-free culture via agarose micromolds

Agarose micromolds were designed and generated as previously described^142^. Each mold consisted of 25 microwells with 500x500x700 µm dimensions and could fit snugly within one well of a 24-well plate (Corning, Corning, NY, #353047) or one well of a 4-well NUNC IVF dish center well dish (Thermo Fisher Scientific, #150260). On the day prior to COC collection and culture, the molds were sterilized by submersion in 70% EtOH for 10 minutes and then autoclaved. 1.5% (w/v) agarose (Hoefer, Bridgewater, MA, #GR140-500) was prepared in DPBS-/- (Thermo Fisher Scientific, #14-190-250), then heated to boiling to dissolve the agarose. About 600 µL of the molten 1.5% agarose was pipetted into the molds, and then the molds were allowed to set for several minutes as the agarose cooled. Solidified agarose micromolds were then placed in 24-well plates or 4-well NUNC IVF dishes and equilibrated overnight in α-MEM/FBS/HEPES/Pyr/mEGF. The next day, COCs were collected as described above and cultured with or without 10 µM PCI. COCs were cultured for 0 h, 4 h, 8 h, or 12 h before capping the mold with molten 1.5% agarose.

Capped agarose micromolds were fixed in Modified Davidson’s Fixative (Electron Microscopy Sciences, Hatfield, PA, #64133-50) overnight, then washed 3x10 minutes in 70% (v/v) EtOH. Fixed molds were processed by an automated tissue processor (Leica Biosystems, Buffalo Grove, IL) and embedded in paraffin wax. Tissue blocks were sectioned at 5 µm thickness. Slides containing COC micromold sections were stained with hematoxylin and eosin using the Leica Autostained XL and following a standard protocol. Stained COCs were imaged at 40x on the EVOS FL Auto imaging system. Magnified images of the matrix at 100x were imaged on the Nikon Eclipse E600. In images where the oocyte was visible, the total numbers of cumulus cells in a single section were counted for control and PCI-treated COCs using Fiji.

### Hyaluronic acid binding protein assay

To evaluate hyaluronan abundance in PCI-treated COCs, we utilized a biotinylated hyaluronic acid binding protein (bHABP; Sigma-Aldrich, #385911). Sections containing control or PCI-treated COCs within COC micromolds were deparaffinized in Citrisolv and rehydrated in decreasing concentrations of ethanol. Slides were washed in 1x PBS. An avidin/biotin blocking kit (Vector Laboratories, Newark, CA, #PK-6101) was used to block any endogenous avidin or biotin on the samples. Avidin and biotin blocks were added to all slides for 15 min each at room temperature (1x PBS washes after each block). Samples were then incubated with goat serum for 20 min at room temperature followed by washing with 1x PBS. After blocking, bHABP diluted in goat serum at 1:100 concentration was applied to all sections. Samples were incubated with bHABP for 1 h at room temperature, then washed in 1x PBS. The Vectastain Elite ABC Kit (Vector Laboratories) and TSA (Tyramide Signal Amplification) plus fluorescein kit (1:400; Akoya Biosciences, Malborough, MA, # NEL741001KT) were used to amplify the signal (1x PBS washes after each amplification step). Slides were mounted in Vectashield Plus Antifade Mounting Medium with DAPI (Vector Laboratories, Newark, CA, #H-2000-10). Three independent trials were performed.

Fiji was used to quantify the HABP signal. COC area was analyzed by outlining COCs with the freehand selection tool and measuring the area of the selected region. Hyaluronan abundance (HABP staining) was analyzed by measuring the thresholded signal of green fluorescence within the region selected for COC area. Thresholds were kept consistent across all COCs within a trial. HABP signal was then normalized by the COC area.

### Live imaging of COC expansion with fluorescent dyes

#### Imaging parameters

COCs were placed in 10 µL maturation media supplemented with SPY650-DNA (1:5000) in µ-Slide 8/18 Well Glass Bottom chambers (Ibidi) at 37°C and 5% CO_2_ in a stage-top incubator integrated with the microscope. Imaging was performed on a laser-scanning confocal microscope (Leica SP8 with HC PL APO 20x CS2 0.75 NA objective and highly sensitive HyD detectors). Samples were excited using a 638-nm laser line, and images were acquired with Nyquist-optimized sampling. Three-dimensional confocal image stacks were acquired at ∼10-minute intervals for up to 16 h. Z-stacks spanned ∼110 μm with 80-90 optical sections (z-step size 1.22 μm), with sampling parameters adjusted slightly between experiments as needed.

#### Cumulus cell tracking

Cumulus cell motility was quantified from 3D time-lapse confocal z-stacks acquired at ∼10-minute intervals over 14 h. Individual cumulus cells were tracked using the Spots module in Imaris (Bitplane, version 9.7.0). Tracks shorter than 6.5 h were excluded from analysis. All trajectories and related parameters were exported and analyzed in MATLAB (MathWorks, R2025b). Mean square displacement ^116,117^ was calculated for each track as:

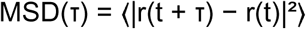

where τ is the lag time, r(t) is the 3D position vector at time t, and the average is taken over all valid time intervals of length τ within each track. Ensemble MSD was computed by averaging individual track MSDs within each experimental group. To account for differences in track number across replicates, MSD was first averaged within each replicate and then across replicates, with SEM calculated at the replicate level.

The anomalous diffusion exponent α was determined by fitting a power law to the ensemble MSD:

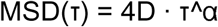

Fitting was performed in log-log space using linear regression over the first 25% of lag times. Per track α values were also calculated by fitting each track independently to assess population heterogeneity. Differences in α between control and PCI-treated groups were evaluated using a permutation test. The observed difference in mean per-track α was compared to a null distribution generated by randomly permuting group labels 10,000 times and recalculating the mean difference. The p-value was defined as the proportion of permuted differences equal to or greater than the observed value. This non-parametric approach avoids assumptions of normality given the broad distribution of per-track α values.

To assess temporal dynamics, a sliding window analysis was performed using windows of 10 consecutive time points (∼1.8 h) advanced in single-frame steps. Within each window, MSD and α were calculated as described above using only positions within that interval. Mean α was calculated across tracks within each group, with SEM across tracks. To align time points across replicates with slightly variability in acquisition timing, timestamps were rounded to the nearest frame interval (655 sec) prior to analysis. Between-group differences in α were assessed at each window using permutation testing (1,000 permutations), with p <0.05 considered significant. Divergence time was defined as the first window in which mean α in controls persistently exceeded that in treated cells.

### Scanning electron microscopy

Mouse COCs were collected and cultured *in vitro* as described above for 12 h in control or PCI-treated media. During IVM, Silicon wafers (Ted Pella, Redding, CA, #1601) were coated in 200 µg/mL poly-D-lysine (EMD Millipore, Burlington, MA, #A-003-E) to promote COC adherence to the wafer. After IVM, COCs were transferred directly from the culture media to the poly-D-lysine coated silicon wafers. As much culture media as possible was removed from the wafers without letting the COCs get completely dry. For all following steps, the appropriate solution was added directly to the top of the wafers to immerse the COCs without overflow of the solution beyond the wafer edges, and a stereo microscope was used to ensure COCs remained adhered to the wafer during reagent transfer. COCs were fixed in 2.5% (v/v) EM Grade glutaraldehyde (Electron Microscopy Sciences, Hatfield, PA, #16220) and 2% PFA (Electron Microscopy Science, Hatfield, PA, # 15710) in a 0.1M cacodylate buffer solution pH 7.2 (Electron Microscopy Sciences, Hatfield, PA, #11654) for 30 min, then post-fixed in 1% (w/v) osmium tetroxide (Electron Microscopy Sciences, Hatfield, PA, #19150). COCs were then dehydrated in a graded series of ethanol dilutions and stored in 100% ethanol overnight at 4°C.

The following day, wafers containing adhered COCs were transferred upright to small microporous capsules and underwent a 15-20 min purge cycle in the Critical Point Dryer (Samdri-795, Tousimis, Rockville, MD). After drying, wafers were mounted on SEM aluminum stubs covered in carbon tape (Electron Microscopy Sciences, Hatfield, PA, #77817-05) Using a conductive silver pen, (Electron Microscopy Sciences, Hatfield, PA, #12644-01) a line of silver ink was drawn from the edge of the edge of each stub to the edge of the COC sample. Wafers were then coated in 9 nm osmium (OPC60A Osmium Coater, Filgen, Nagoya, Japan). All images were captured on the Hitachi SU8030 (Tokyo, Japan) at 10 kV. Two independent trials were performed.

### Adhesion assay

Adhesion of cumulus cells was measured using the xCELLigence Real-Time Cell Analyzer (RCTA). The plates used in this system contain microelectronic sensors to correlate electrical impedance with cell adhesion to an ECM substrate^143^. Here, purified human fibronectin (BD Biosciences) was utilized as the ECM substrate, as it is a known component of the ovarian follicle wall^144–146^. Wells of the xCELLigence RCTA electronic plate (E-plate 16) were coated with fibronectin for at least 4 h at 37°C and 5% CO_2_, before washing with PBS. Wells were then preincubated with α-MEM and 250 µM pyruvate at 37°C and 5% CO_2_.

This assay utilized cumulus cells from postovulatory (mature) COCs. To collect postovulatory COCs, mice were hyperstimulated with an IP injection of 5 IU equine chorionic gonadotropin (eCG) to recruit follicles. 46 h after eCG, mice were induced to ovulate with 5 IU hCG. COCs were collected from the ampulla of the oviduct at 14 h post-hCG injection, and 10 COCs were added to each well. Adhesion was tracked for at least 12 h with measurements taken every 3 min. The following calculation was used to evaluate the adhesion index: R_n_-R_b_ (R_n_ = cell-electrode impedance of the well in the presence of cells; R_b_ = background impedance of the well containing only media).

### Quantification and statistical analysis

#### Statistical analysis

Each experiment presented in this study was repeated at least two times. Each COC was considered an individual biological replicate unless otherwise specified in figure legends. Graphpad Prism was used to graph results. Data within graphs are shown as the mean ± the standard error of the mean unless otherwise specified in figure legends. For non-categorical variables, the Shapiro-Wilk test was used to determine normality. Student’s t-tests were used to compare means between two groups. One-way or two-way analysis of variance (ANOVA) tests were used to analyze multiple comparisons for one or two independent variables, respectively. Mixed-effects analysis was used in place of ANOVA for unbalanced data. Tukey’s honest significance test was used to correct for multiple comparisons. The Wilcoxon signed-rank test was applied for analyses of the bulk RNAseq data, and t-tests with Storey’s method was utilized for analyses of the proteomics data. For categorical variables, Fisher’s exact test was used. P-values of less than 0.05 were considered statistically significant.

### Key resources table

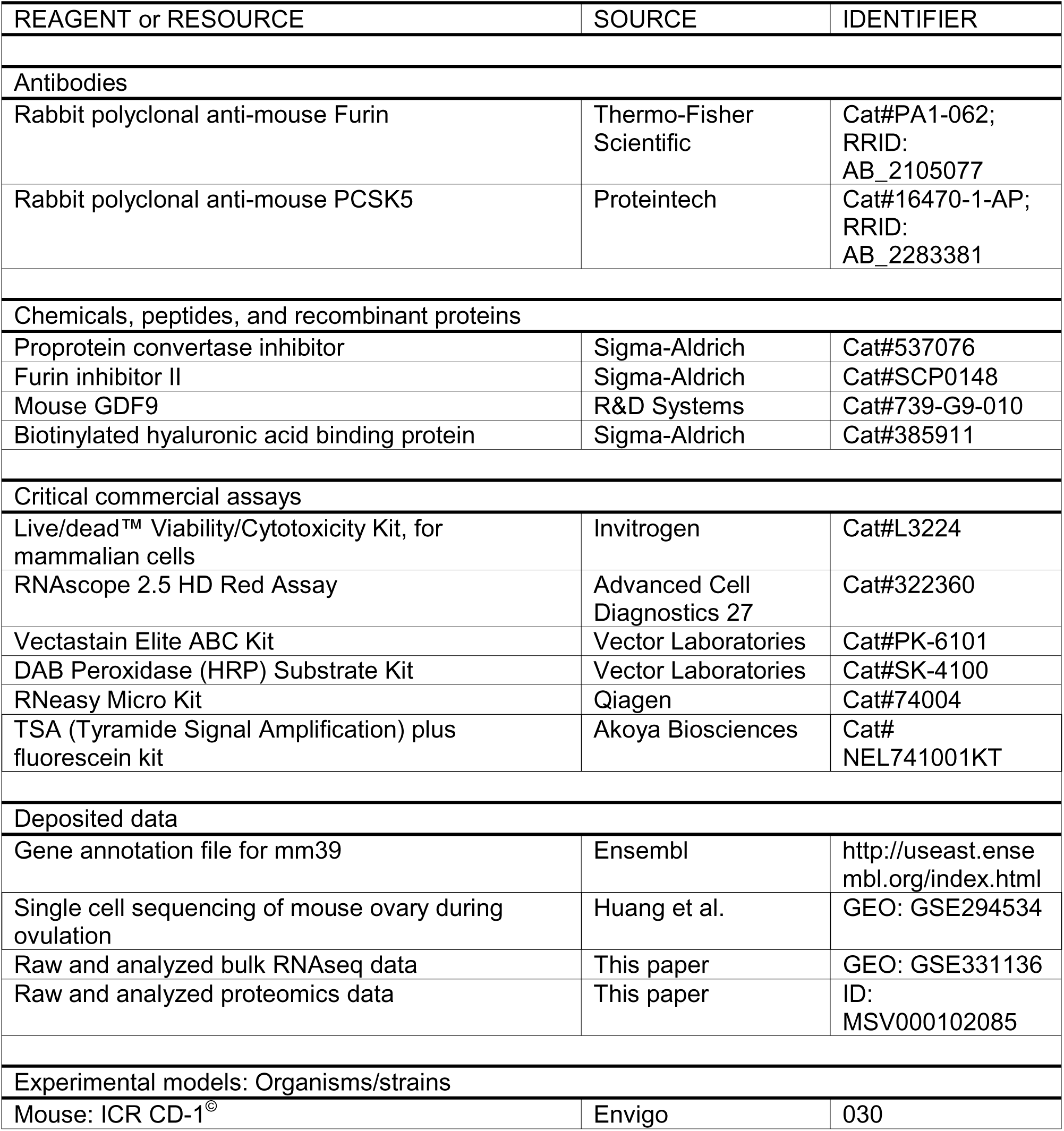
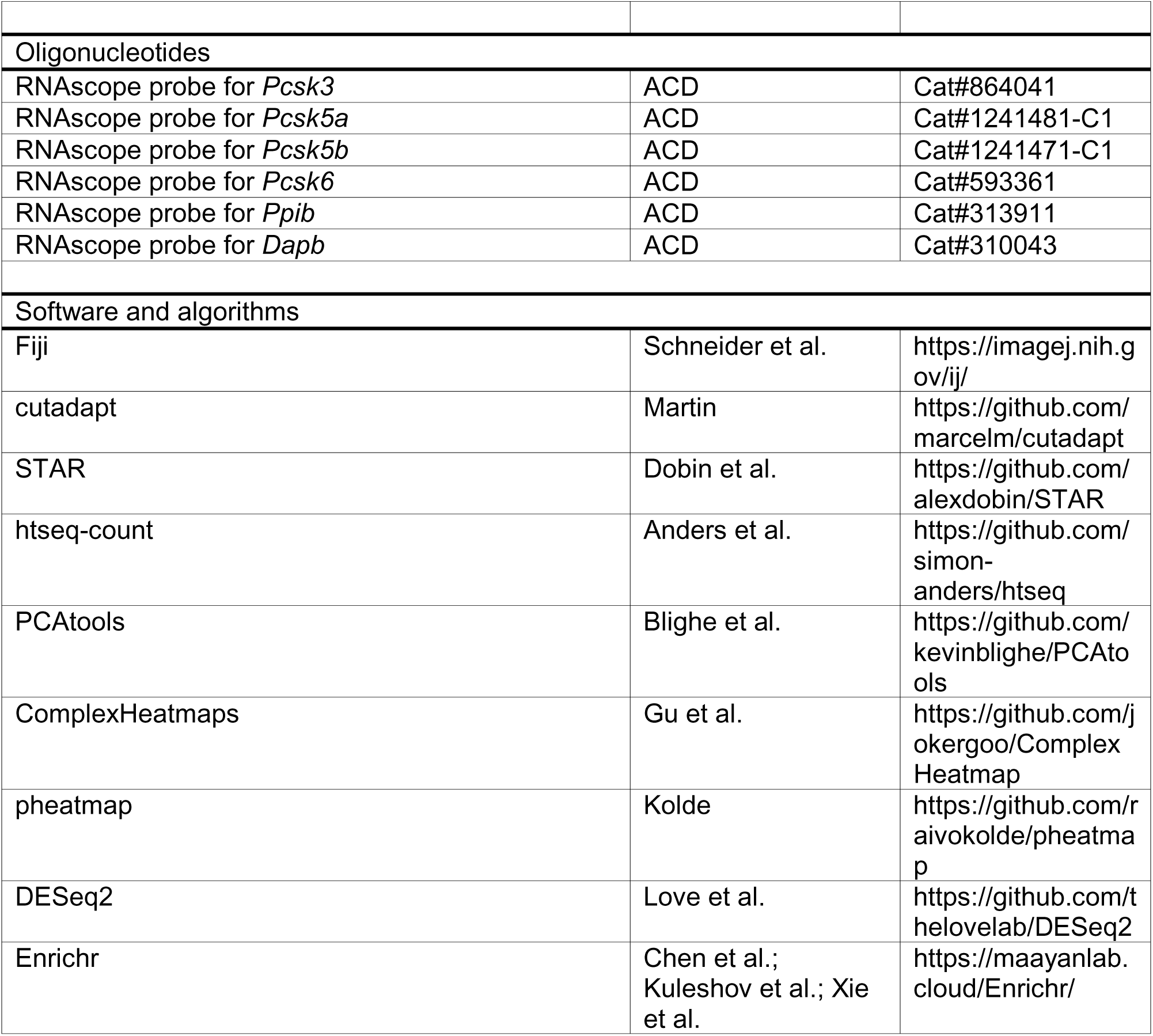

