## Supplemental Figures for "Proprotein convertase activity regulates cumulus-oocyte-complex matrix integrity and cumulus cell migration during ovulation via a GDF9-dependent mechanism"

**Supplemental Figure 1**

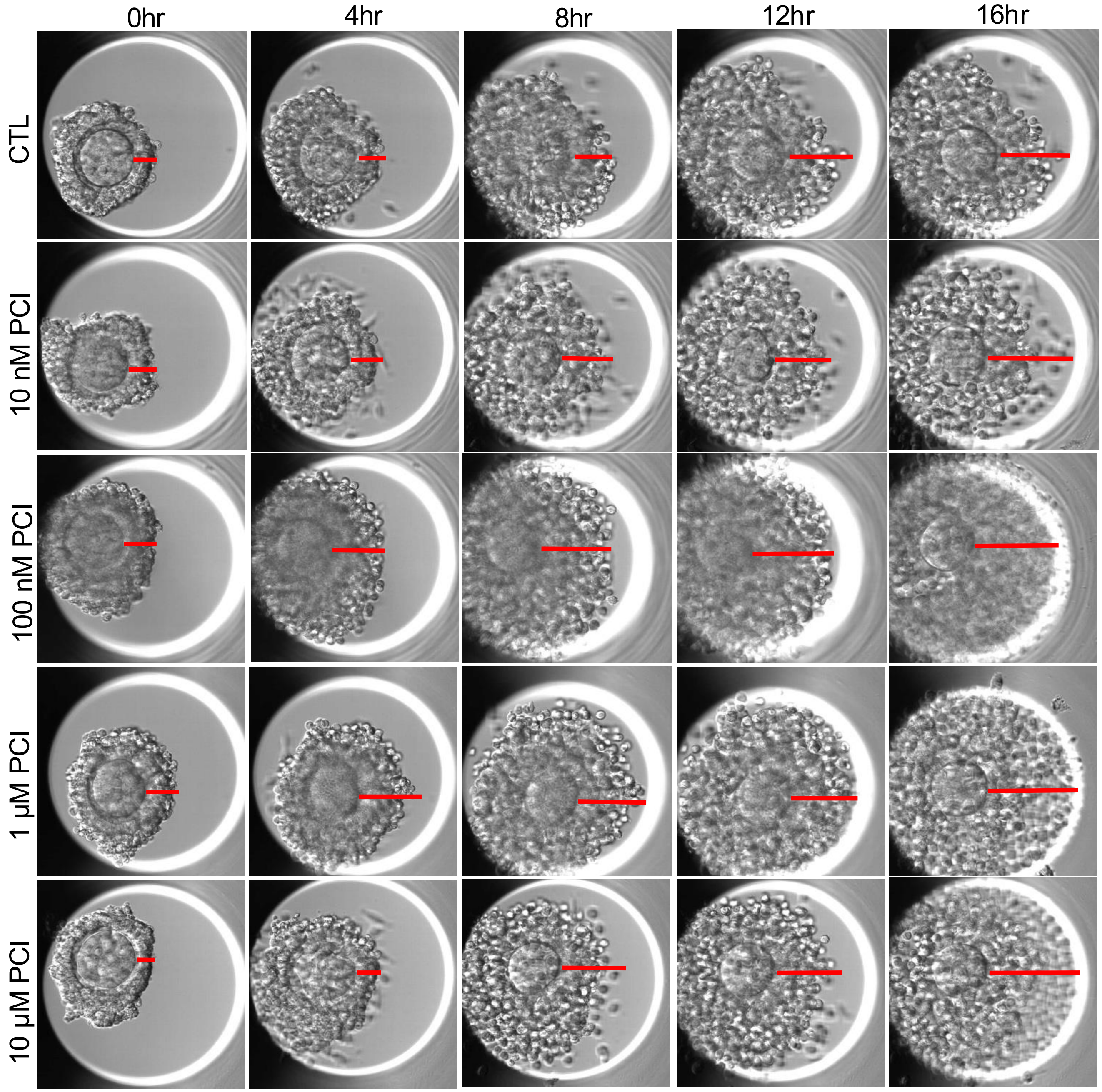

Supplemental Figure 2

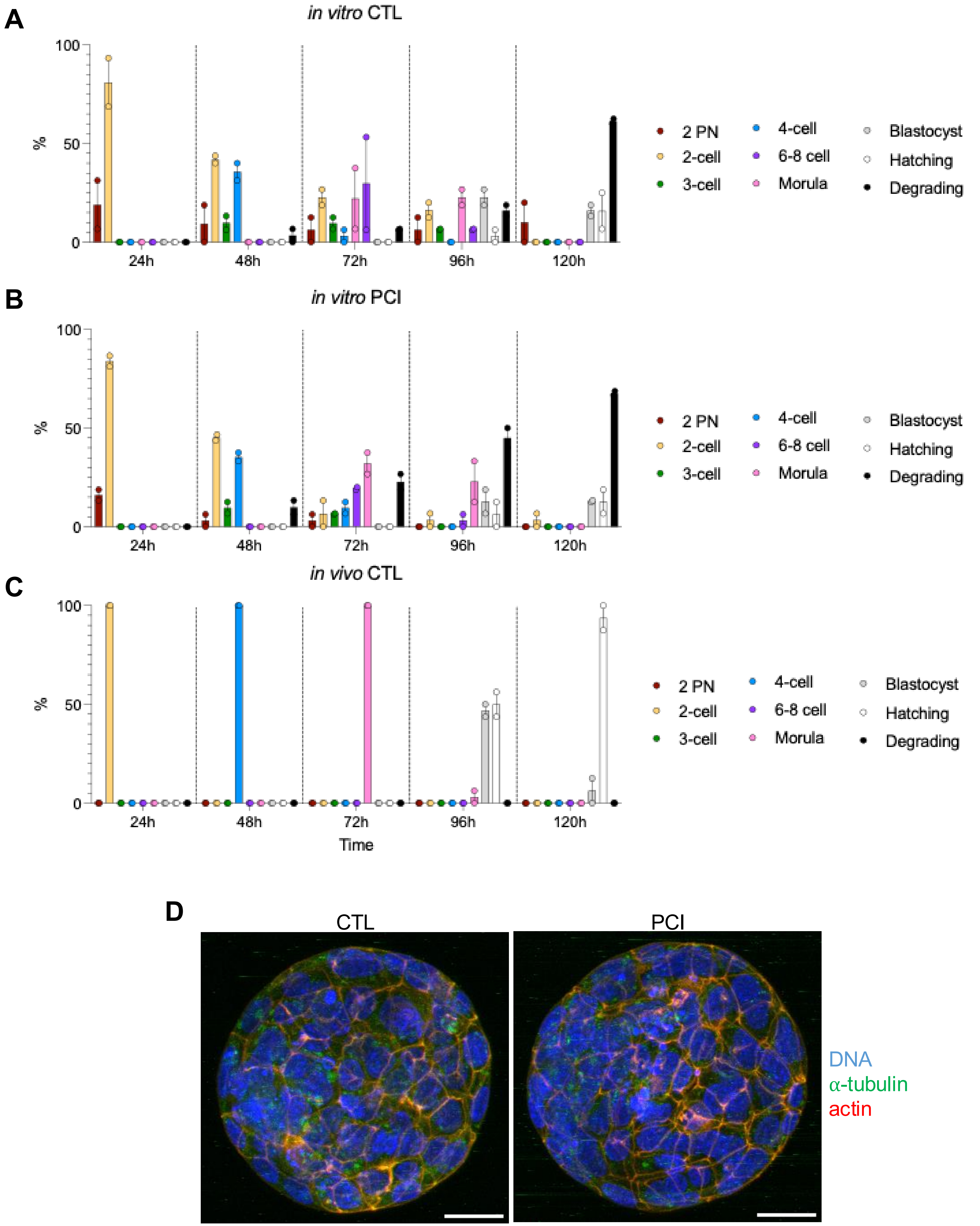

**Supplemental Figure 3**

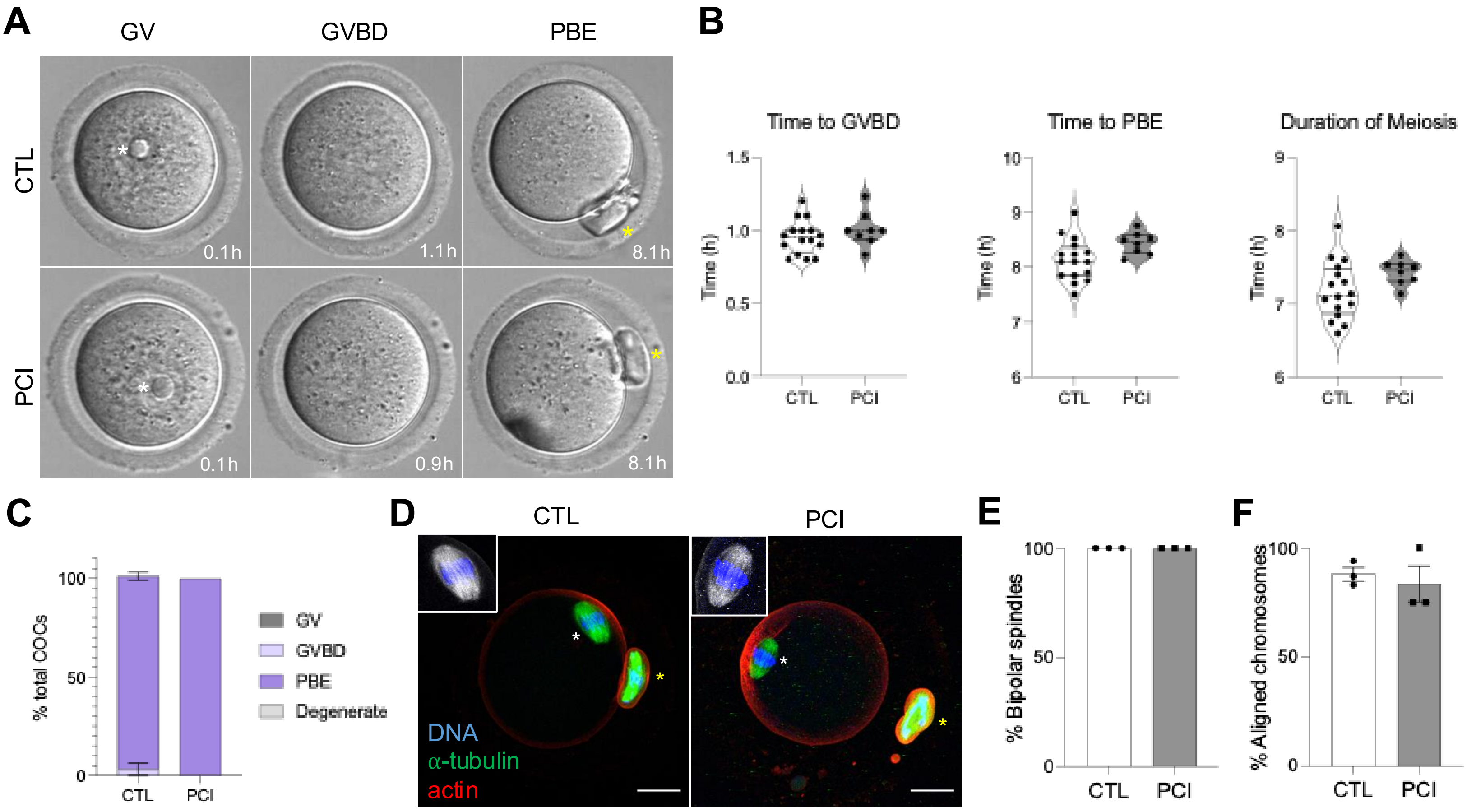

**Supplemental Figure 4**

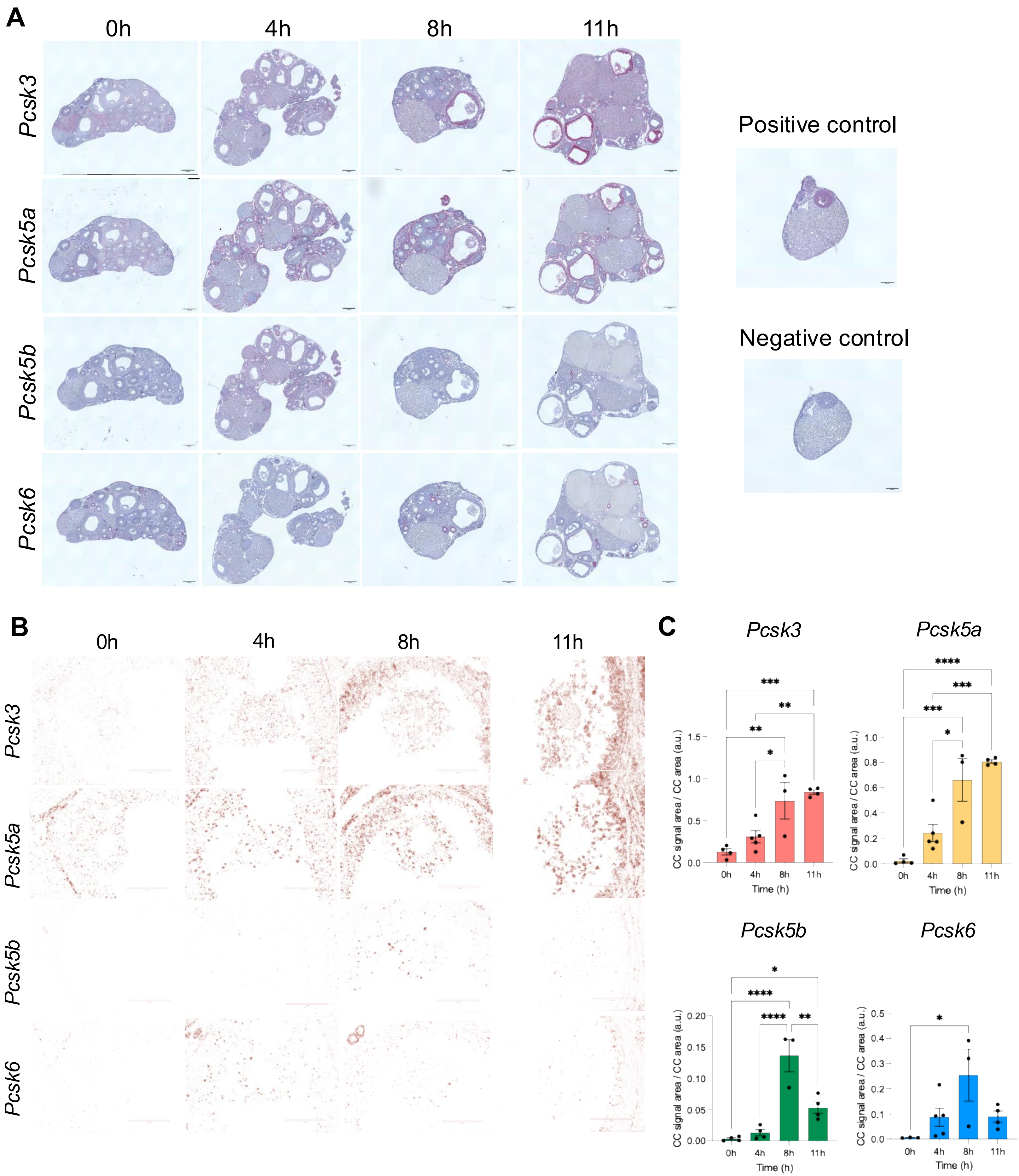

**Supplemental Figure 5**

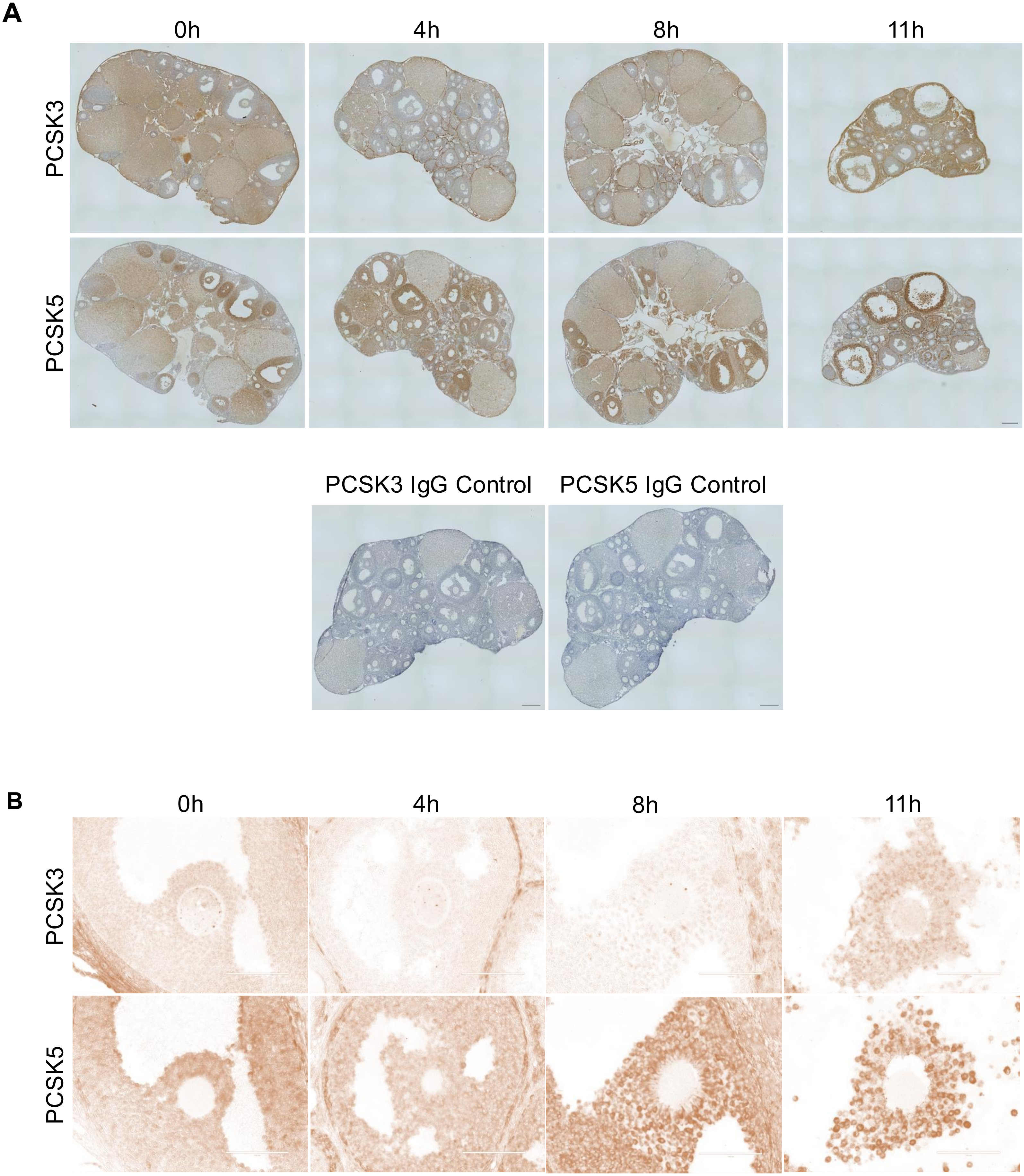

Supplemental Figure 6

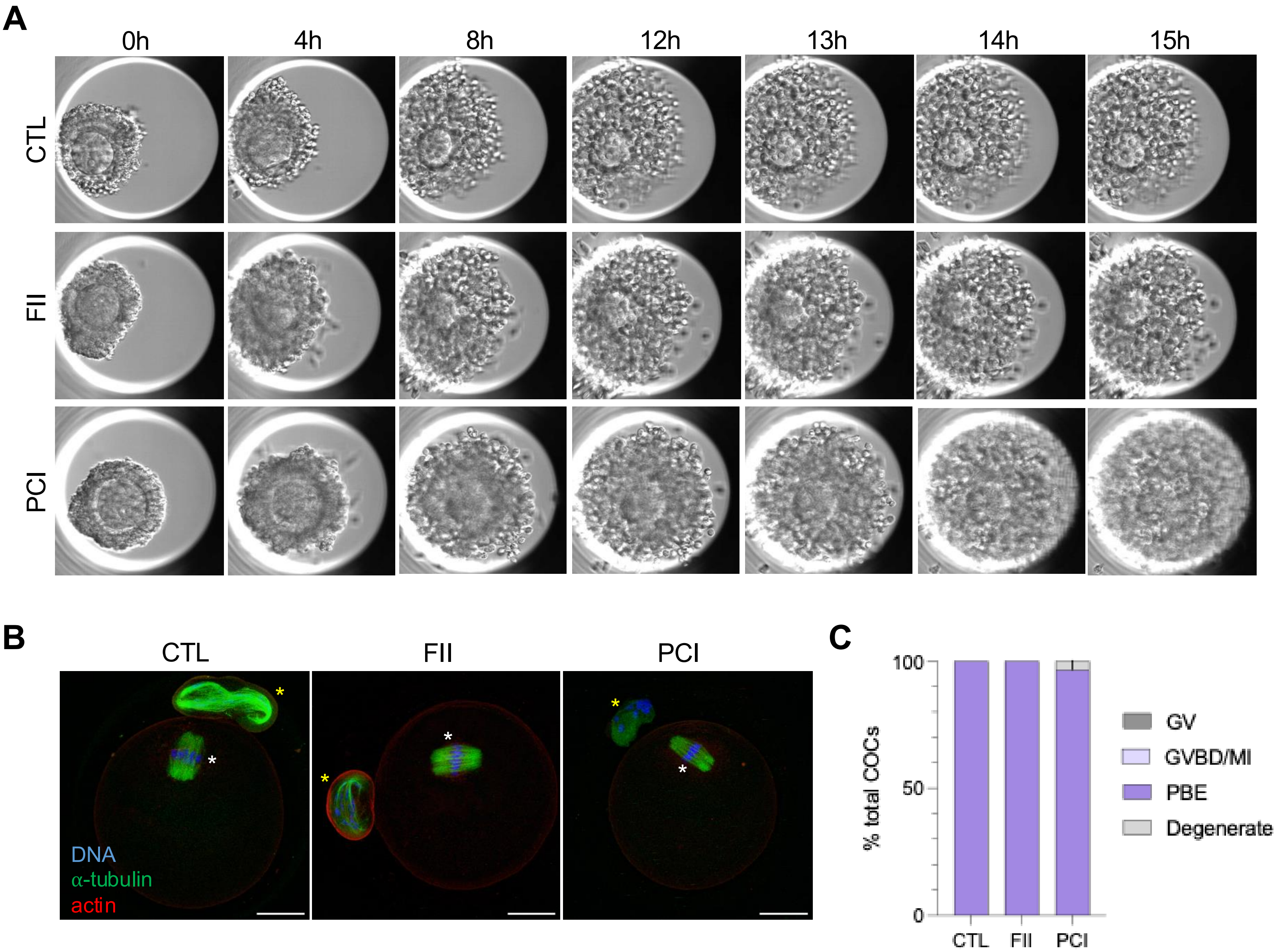

**Supplemental figure 7**

A

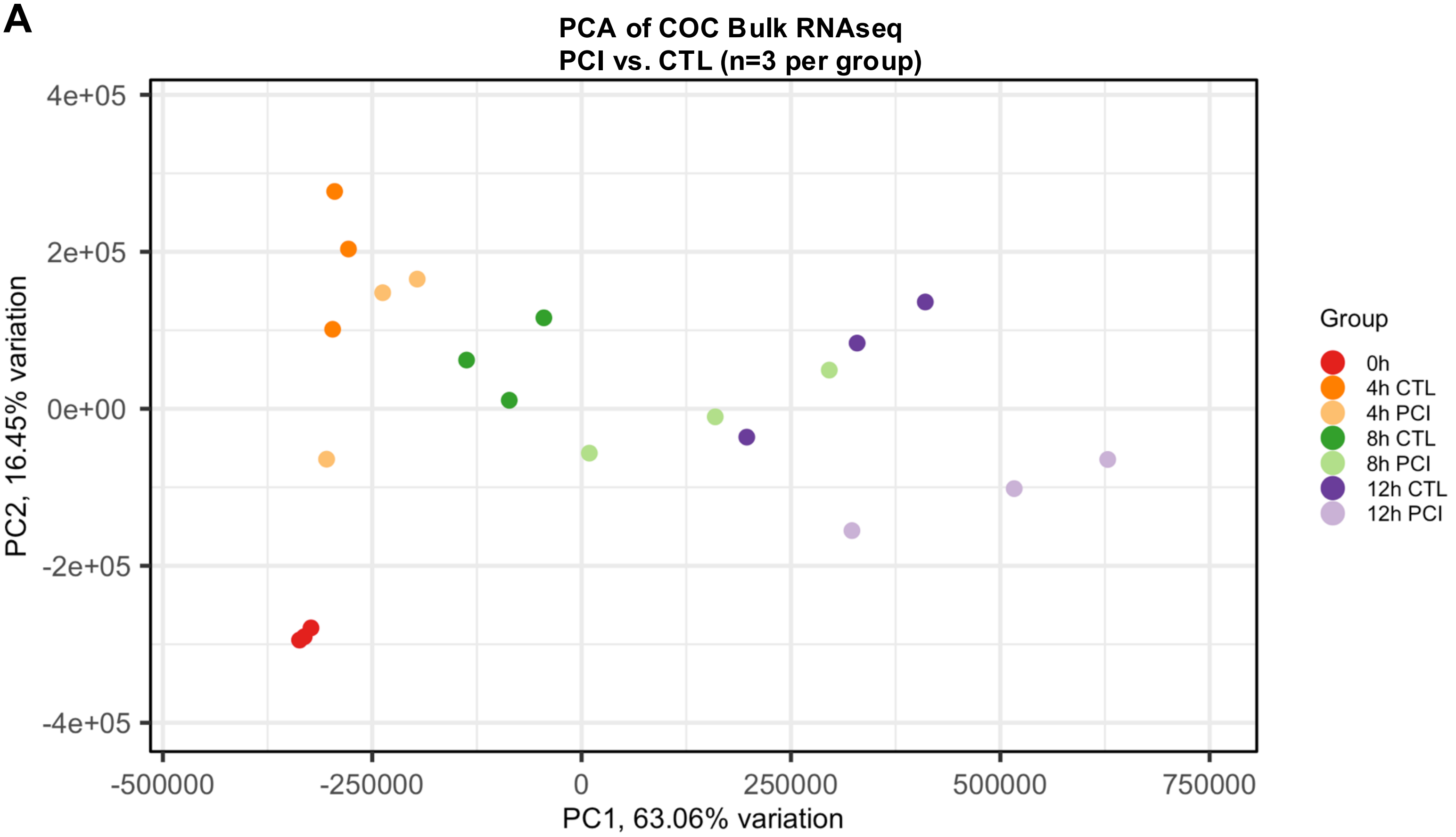


B

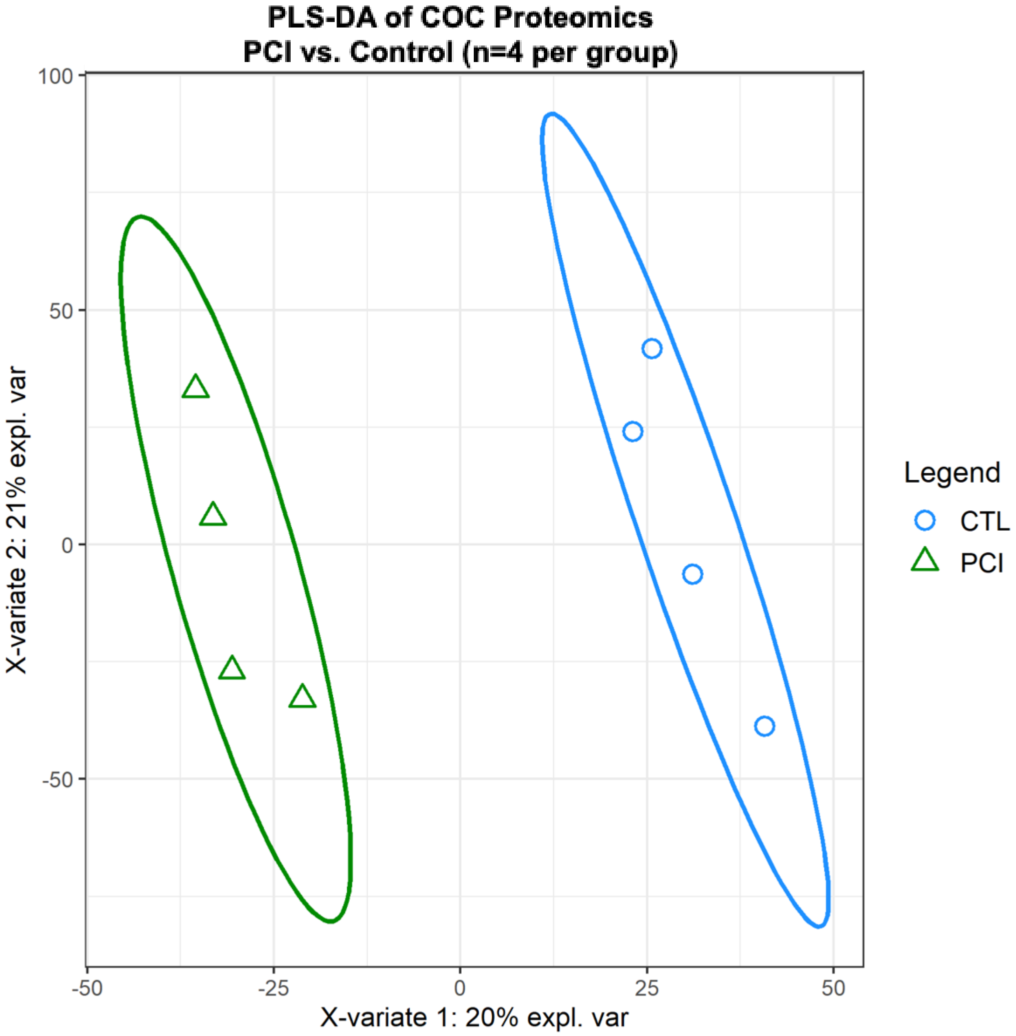

**Supplemental Figure 8**

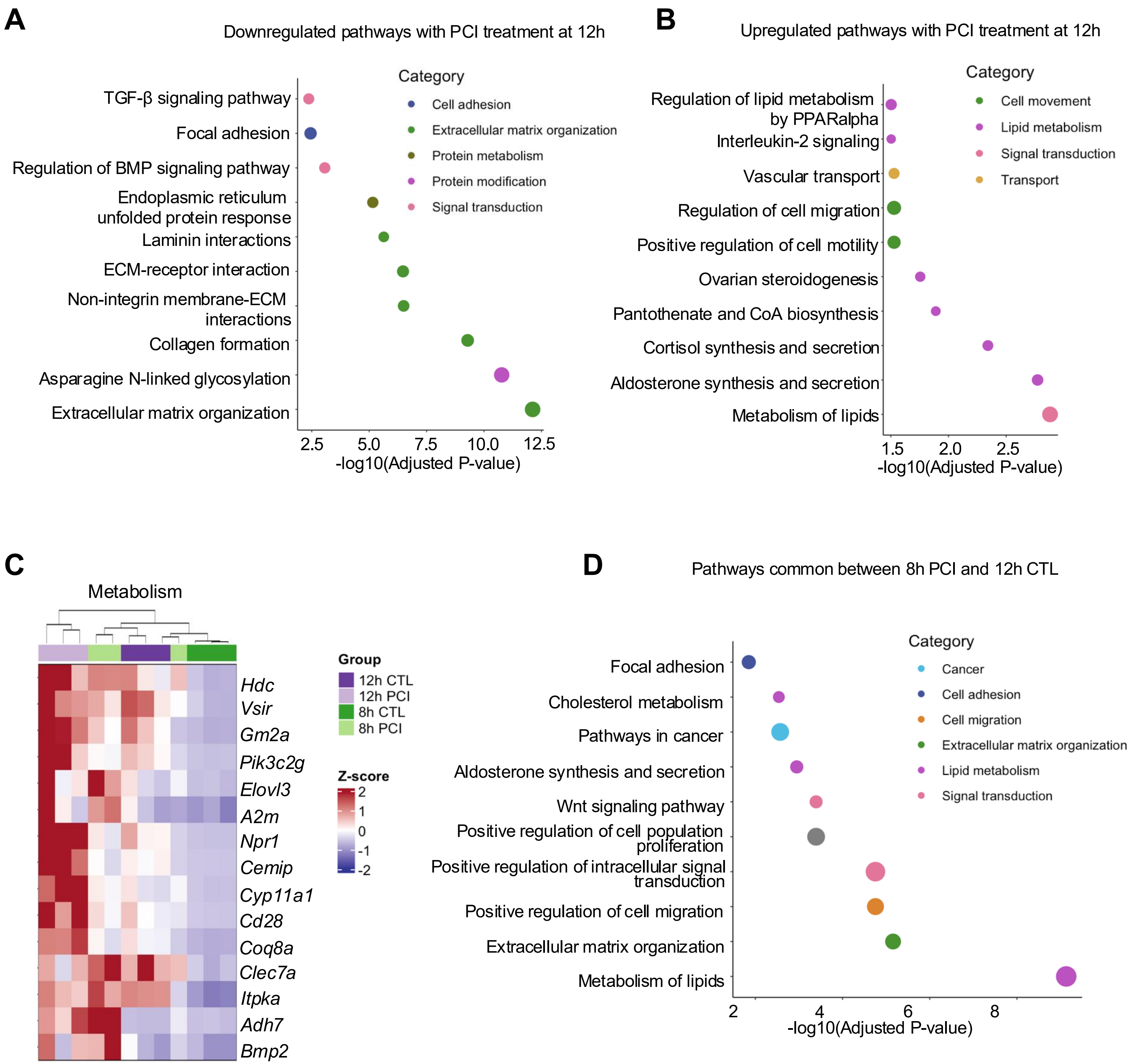

**Supplemental Figure 9**

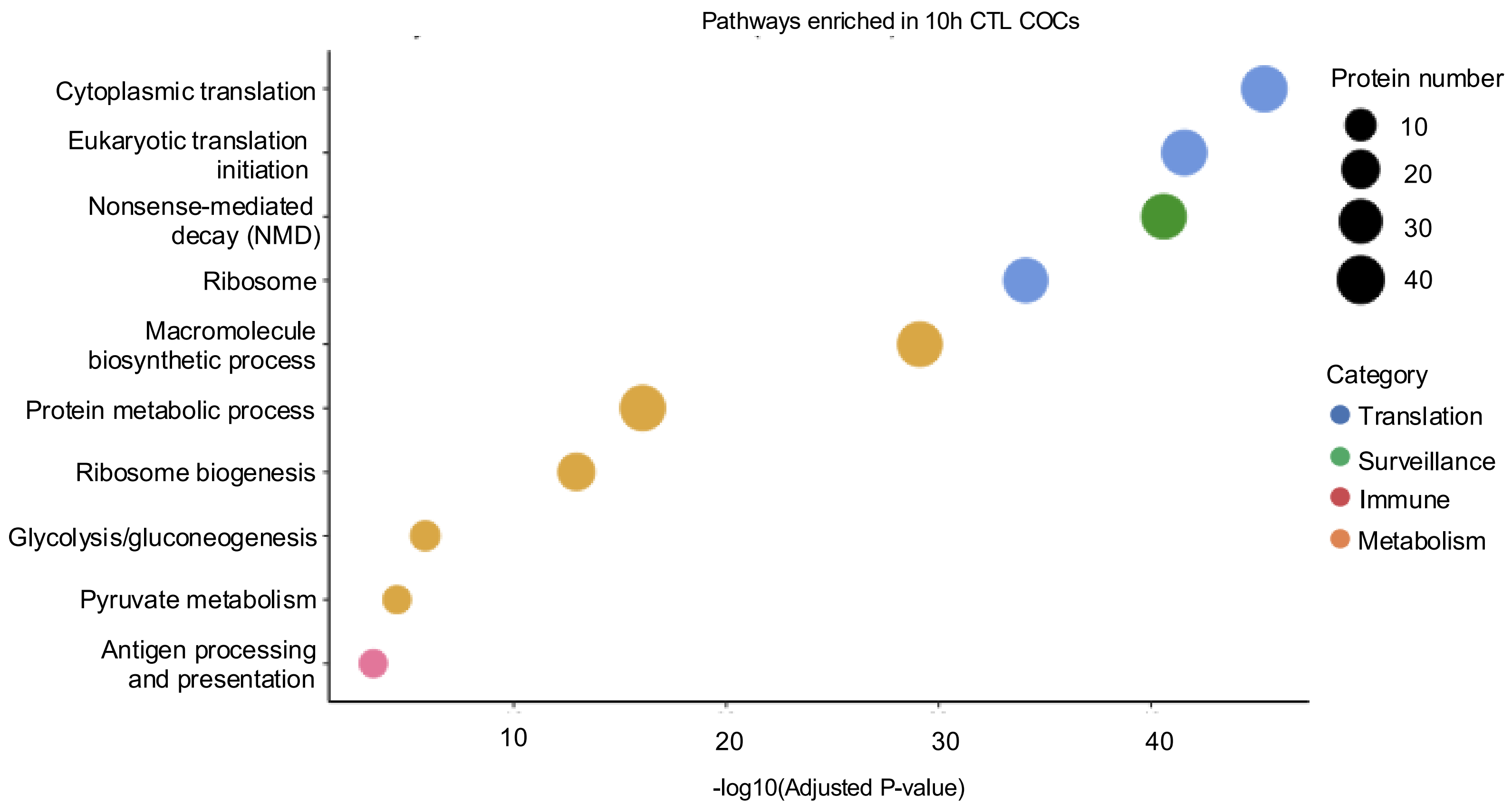

**Supplemental Figure 10**

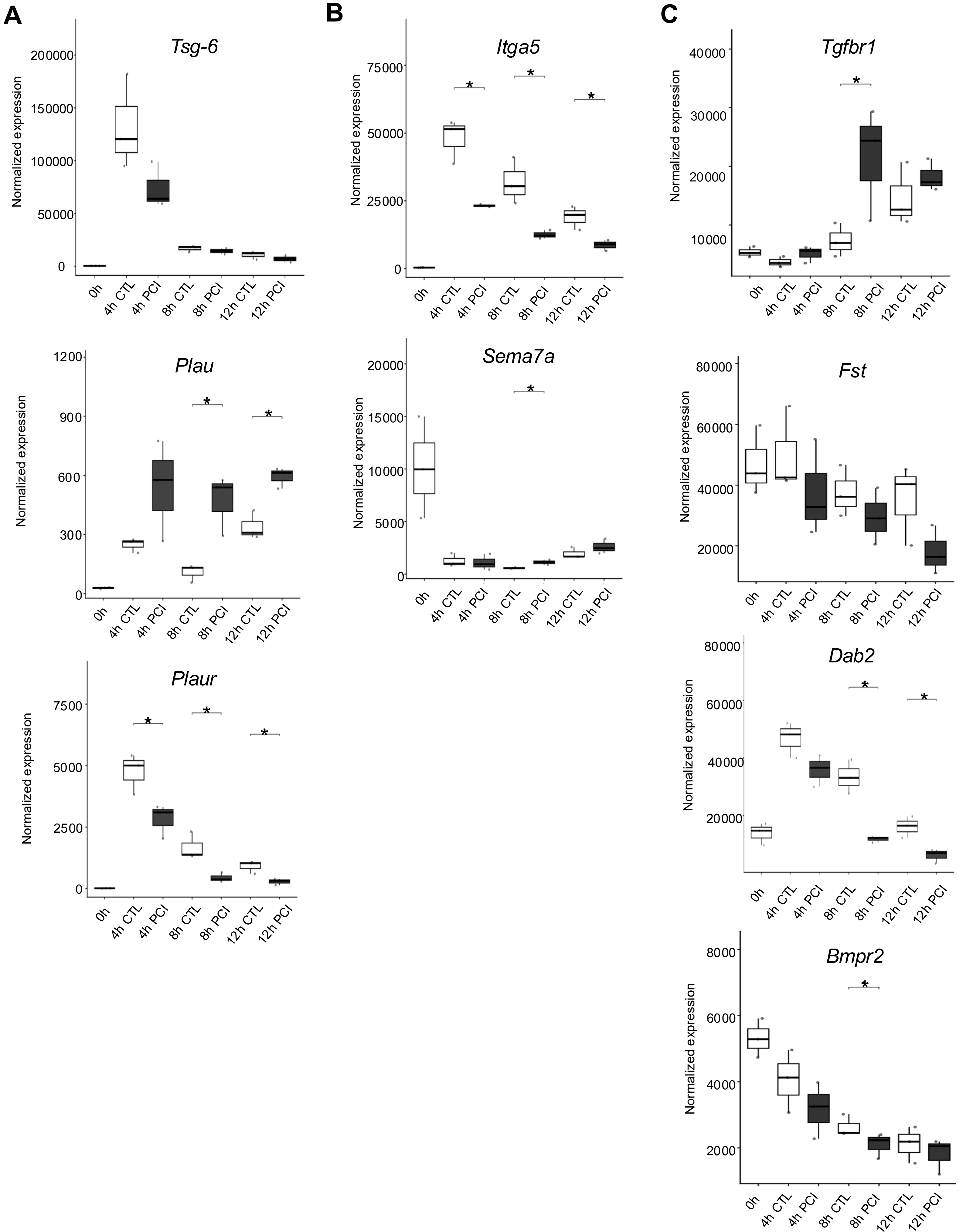

**Supplemental figure 11**

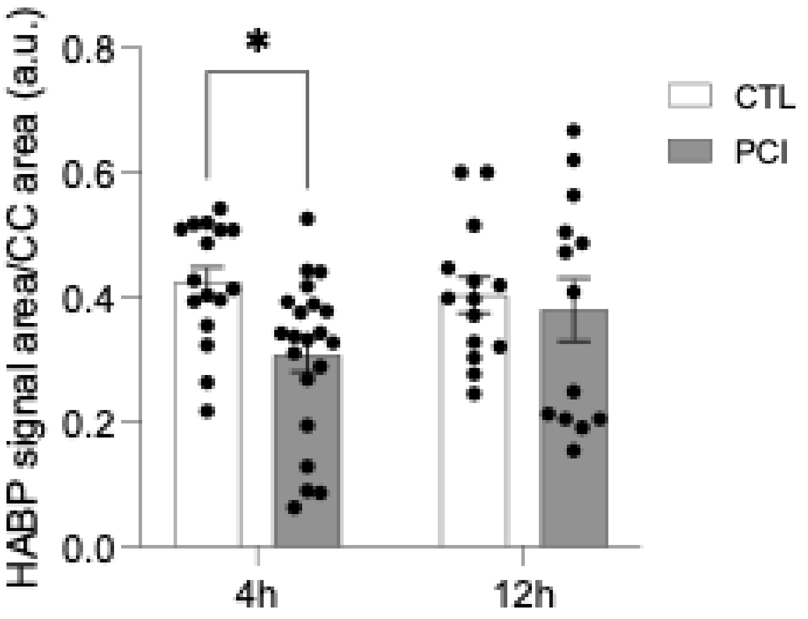

**Supplemental figure 12**

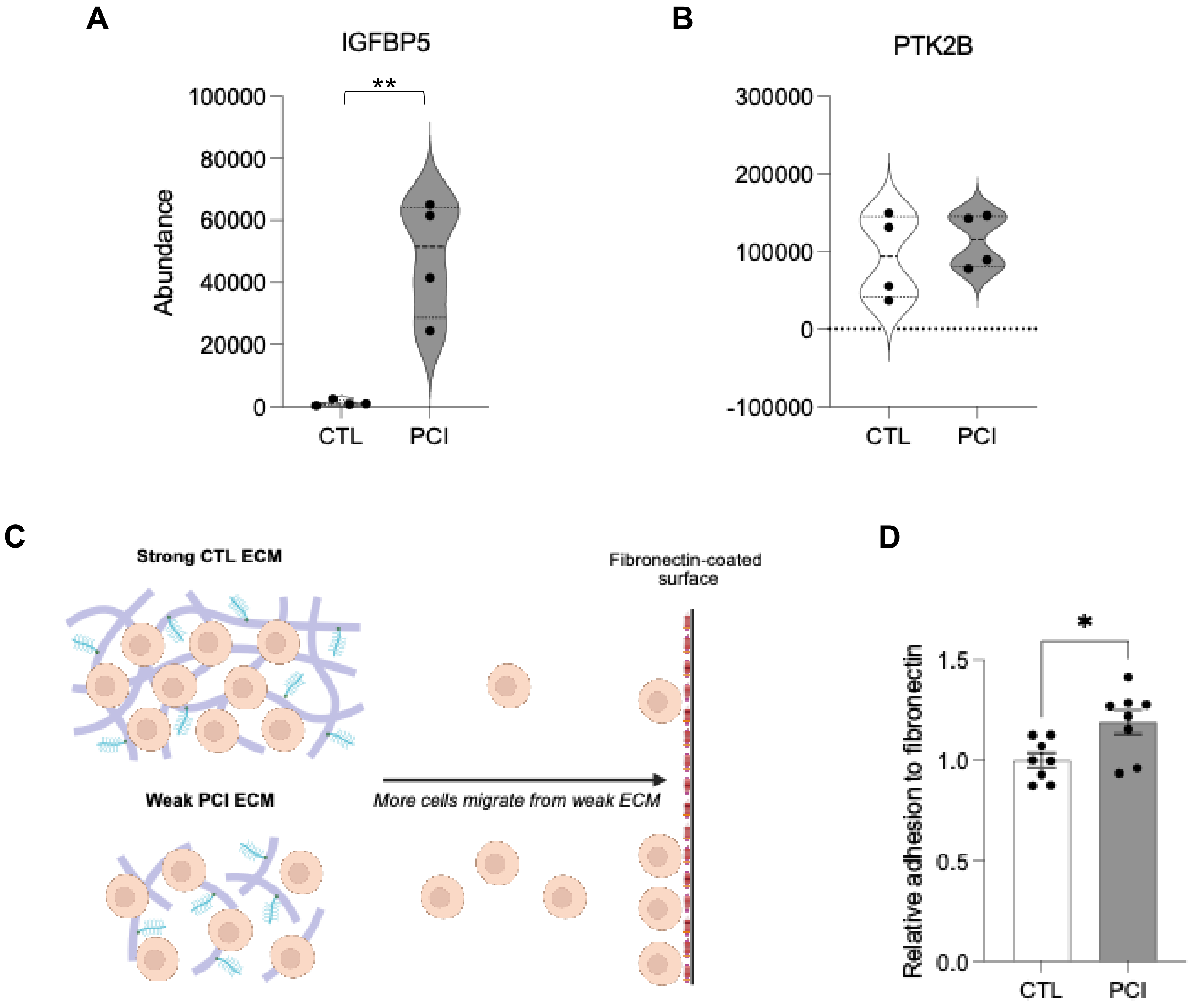

**Supplemental Table 1**

| Category | Genes |
|---|---|
| ECM organization and cell adhesion | ITGB5; LAMB3; LAMA1; LAMA4; TNC; DMP1; LAMC2; THBS1; COL2A1; IBSP; ITGA11; COL6A3; ITGA5; VAV3; PDGFRA; JUN; CCND2; FLNC; PAK3; RAF1; FGF2; LAMA5; ITGB3; ITGA2; ITGA1; COL1A1; COL4A2; COL4A1; DAG1; SDC1; ITGAV; CD36; CD44; RASGRF1; IGF1; CAPN2; TGFB1; COL11A1; NTN4; COL18A1; COL7A1; NID1; COL16A1; PCOLCE2; COL12A1; FBLN2; SCUBE3; NCAM1; COL27A1; DST; LAMB3; MMP2; MMP12; MMP13; BMP1; LOX; P4HA2; ADAM12; ADAMTS10; FLRT2; ADAMTS19; CREB3L1; SERPINH1; HAS2; GSN; APLP1; MMP11; SMOC2; MMP14; P4HA1; P4HA3; CTSL; JAM2; P3H1; BMP4; MFAP5; MFAP2; P4HB; PLEC |
| Cell movement | RET; CSF1R; IGSF8; GRN; CEMIP; SEMA7A; MCTP1; PLXND1; CITED2; SEMA3B; SEMA3G; ARID4A; RORA; PTN; VSIR; EGFR; ROBO1; PPP3CA; CLEC7A; PLAU; PDGFD; GNA12; CYP1B1; FLNA; PLXNA1; PTK2B; PHACTR1; PLXNA3; CARMIL1; ANXA1; IGFBP5; FZD4; SEMA4B; INSR; CAV1; PRKCA; PPP2R3A; F3; TGFBR1; TGFBR3; BMP2; SFRP2; MYADM; SGK1; FERMT2; VCLCAVIN1; SCARB1; SPARC; SYDE1; CLASP2; FN1; GNPDA1; RIN2GRB7; SEMA3D; SEMA3A; CX3CL1; GLUL; PLXNA4; RDX; FRMD5; CXCL12; KIF26A; ANGPT1; WNT5A; PRKD1 |
| Metabolism | RAMP2; CALCOCO1; RAMP3; DDX3X; TFRC; CITED2; IRS1; HFE; NTS; AREG; VSIR; ROBO1; C1QTNF1; HMGN5; CLEC7A; MYC; UBR5; RIPK1; SLC38A2; EGR1; MEF2C; INSR; FN1; F3; TGFBR1; TGFBR3; P2RX7; SFRP4; BMP2; RGCC; SLC6A9; CD28; LIMS1; BMPR1A; NFE2L2; ABCA1; SCARB1; MYLIP; LRP1; LPL; LIPA; SOAT1; STAR; VAPA; NPC2; ANGPTL4; LDLRAP1; LDLR; NPR1; HSD3B1; ITPR2; PRKCA; ATP1B2; ATP1A1; ATP1B1; NR4A2; LIPE; NR4A1; CREB3; PLCB3; PRKD3; CYP11A1; CAMK2G; KL; RUNX2; EGFR; PTHLH; LRP6; SLC9A3R1; MMP16; GNA12; FADS2; SCP2; ELOVL5; ELOVL3; ELOVL6; HACD3; FADS1; ACAA2; PPT1; HADH; MRAP; SLC44A3; HEXB;I NPPL1; RORA; PIK3C2G; AHR; GM2A; FDXR; ME1; CYP1B1; HMGCS2; NCOA2; PCYT1A; DGAT1; ACSL4; SUMF1; ACLY; CYP2U1; PITPNM1; FDX1; PLBD1; AHRR; FAR1; SLC27A3; PLPP2; IDI1; SLC22A5; SGMS2; ACAT2; SGPL1; ALDH3B1; PSAP; SC5D; SCAP; SPTSSA; TBL1X; HAO2; OSBPL9; CYB5B; CTSA; B3GALNT1; BCHE; CHKA; OSBPL3; CAV1; MBOAT1; PTPN13; ACSF2; SGPP1; PIKFYVE; GPAM; ETNK2; LPCAT3; DHCR7; RGL1; GLA; LGMN; PANK1; SAT1; HS6ST1; NDST2; NAGLU; PDK4; ENPP1; NUDT13; CSGALNACT2; SLC6A12; HSPG2; RBP4; BTD; CEMIP; ISYNA1; MAOA; FUT11; MGST1; GMPR; IQGAP1; ADH7; PRKAR2B; HMOX1; IP6K3; PCK2; GSTM1; GOT1; IDH1; PAICS; QDPR; GCLC; G6PC3; VNN1; AKR1B10; TCN2; KYAT3; NMT2; SERINC3; HCCS; OPLAH; NAPRT; ME2; HIBCH; TPI1; ABCC5; SORD; CTPS2; PPP2R5D; ITPKB; VCAN; TST; IVD; DPYD; ITPKA; ALDH1A1; BLVRB; R; BCAT1; ASAH1; INPP1; HDC; ODC1; PPM1K; PAPSS1; UGP2; GPC4; SLC36A4; XDH; CBR3; NQO2; HS3ST1; PDP1; GNPDA1; COQ8A; PFKP; V; PCSK5; A2M; ANGPT1; RDX; WNT5A; OLFM1; DNAJA4; ID2; TLR3; MARK1; PRKCB; ITPR1; PRKD1; CACNA1H; KCNK3; VNN3; FSHR; HSD17B7; UGCG; GAL3ST1; ENTPD1; PPAT; PDE3A; PDE4B; PDE5A; AMPD3; INSIG1; NPAS2; ARNTL; ENPP6; ARNT2; GK; SRD5A1; LPIN3; TK2; GLUL; AASS; HGSNAT ;GLCE; CACNB2; CARNMT1; SHMT1; LDHB; BRIP1; SULT1E1; PRODH; HPSE; ALAS1 |

**Supplemental Table 2**

| Oocyte marker | Average $Log_2$ Ratio | Q-value |
|---|---|---|
| BMP15 | 0.71 | 0.180 |
| GDF9 | 0.25 | 0.659 |
| ZP2 | 0.08 | 0.367 |
| DDX4 | -0.14 | 0.725 |
| NPM2 | 0.33 | 0.00841 |
| ZAR1 | 0.12 | 0.373 |
| YBX2 | 0.33 | 0.785 |
| DPPA3 | 0.04 | 0.407 |
| SUB1 | 0.50 | 0.183 |
| ZP3 | 0.11 | 0.387 |
| NLRP5 | 0.25 | 0.00599 |
| ZP1 | 0.28 | 0.219 |
| KIT | 0.47 | 0.418 |
